# Structural basis of NSD2 degradation via targeted recruitment of SCF-FBXO22

**DOI:** 10.1101/2025.08.29.673087

**Authors:** Kevin C. Robertson, Sascha J. Amann, Tongkun Liu, Adam V. Funk, Xianxi Wang, Irina Grishkovskaya, John R. Tabor, Jacqueline L. Norris-Drouin, Cheryl H. Arrowsmith, Jon L. Collins, Michael J. Emanuele, David Haselbach, Lindsey I. James, Nicholas G. Brown

**Author notes:** Correspondence should be addressed to D.H., L.I.J., or N.G.B. These authors contributed equally: K.C.R., S.J.A., and T.L.

## Abstract

Targeted protein degradation (TPD) through the ubiquitin-proteasome system is driven by compound-mediated polyubiquitination of a protein-of-interest by an E3 ubiquitin (Ub) ligase. To date, relatively few E3s have been successfully utilized for TPD and the governing principles of functional ternary complex formation between the E3, degrader, and protein target remain elusive. FBXO22 has recently been harnessed by several groups to target different proteins for degradation. FBXO22 recruitment has been enabled through degraders that covalently modify its cysteine residues. Here, we reveal that the aldehyde derivative of UNC10088 promotes cooperative binding of FBXO22 to NSD2, a histone methyltransferase and oncogenic protein, leading to a cryo-EM structure of the full SKP1-CUL1-F-box (SCF)-FBXO22 complex with NSD2. This structure revealed a conformational change in the FBXO22 loop surrounding C326, further exposing the cysteine for covalent recruitment. Additional medicinal chemistry efforts led to the discovery of benzaldehyde-based non-prodrug degraders that similarly engage C326 of FBXO22 and potently degrade NSD2. Furthermore, unlike many degraders, our molecules recruit NSD2 to a different surface of FBXO22 than the known FBXO22 substrate BACH1, allowing for concurrent complex formation and degradation of both the neosubstrate and endogenous substrates. Overall, we demonstrate the biochemical and structural basis for NSD2 degradation, revealing key principles for efficient and selective TPD by SCF^FBXO22^.

## INTRODUCTION

Targeted protein degradation (TPD) involves the co-opting of cellular protein degradation machinery to destroy disease-causing proteins. Degrader molecules, e.g., molecular glues (MGs) and proteolysis-targeting chimeras (PROTACs), function by bringing an E3 ubiquitin ligase and target protein together to form a ternary complex^1-3^. This induced proximity facilitates polyubiquitination of the target protein, promoting its proteasomal degradation. TPD offers multiple benefits over traditional pharmacological efforts. For example, TPD uses an event-driven, catalytic mechanism, allowing for the compounds to bind anywhere on their target protein, bind less potently than traditional inhibitors, and be used at sub-stoichiometric levels. Therefore, a lower concentration is often needed for the same potential therapeutic benefit^2,4^.

The ubiquitination process requires a set of E1, E2, and E3 enzymes. The E1 activates and facilitates the transfer of Ub to E2s. At this point, the E2∼Ub (∼denotes a thioester intermediate) can cooperate with a RING type E3 and transfer the Ub directly to the substrate or transfer the Ub to a catalytic cysteine in HECT- or RBR-type E3s, which then modify the substrate^5,6^. The human genome encodes >600 E3 Ub ligases. The majority fall under a certain family of E3s known as Cullin-RING ligases (CRLs).

CRLs can be broken down into five broad subfamilies that contain dozens of substrate receptors, which are the targets of the E3 ligands commonly used in PROTACs^7^. To date, most TPD reagents use ligands for either von Hippel-Lindau (VHL) or Cereblon (CRBN) that are assembled with CUL2-RBX1 and CUL4-RBX1, respectively^2^. However, there are hundreds of substrate receptors, representing a vast, largely untapped opportunity for the development of novel approaches to TPD.

The SKP1-CUL1-F-box (SCF) family of Ub ligases uses CUL1-RBX1 as the catalytic scaffold while the ∼70 F-box proteins recruit various substrates. Several recent studies identified that a particular F-box protein, FBXO22, which is overexpressed in several cancers^8^, can be harnessed for TPD of several neosubstrates, including NSD2 (nuclear receptor-binding SET domain-containing 2), XIAP (X-linked inhibitor of apoptosis protein), FKBP12 (FK506-binding protein 12), and SWitch/Sucrose nonfermentable (SWI/SNF) complex ATPases (SMARCA2/A4). We previously identified that our primary amine-containing degraders were being metabolized into an active aldehyde species that forms a reversible covalent bond with FBXO22 on Cys326 to promote NSD2 degradation^9^. These findings were also corroborated in Kagiou et al.^10^ Interestingly, Basu et al. found that an electrophilic α-chloroacetamide group can be employed to engage FBXO22 via two different cysteines (C227, C228) to promote target degradation^11^. A recent preprint (Villemure et al.) also reported a dependency on C228 and C326 for FBXO22 recruitment and the subsequent degradation of SMARCA2/A4. All of the cysteines used to recruit FBXO22 for TPD are on one face of the protein which is different from the surface used to recruit its native substrate BACH1^12,13^. Despite the availability of numerous FBXO22-recruiting degraders, a structure of FBXO22-recruiting a neosubstrate is not currently available.

Our prior work has focused primarily on targeted degradation of the methyltransferase and methyl-lysine binding protein NSD2, which is an oncogene and a high value therapeutic target in several cancers including multiple myeloma, acute lymphocytic leukemia, and prostate cancer^14-18^. We have shown that our previously reported potent and selective NSD2 degrader, UNC8732, is metabolized enzymatically and the corresponding aldehyde species effectively recruits FBXO22 to promote ternary complex formation and subsequent degradation of NSD2 (Figure 1A). We also reported an analogous compound, UNC10088, in which the amine of UNC8732 is replaced with a sodium bisulfite adduct, allowing for conversion to the active aldehyde species via simple hydrolysis. UNC10088 is equally as effective as UNC8732 in degrading NSD2.

**Fig. 1:**
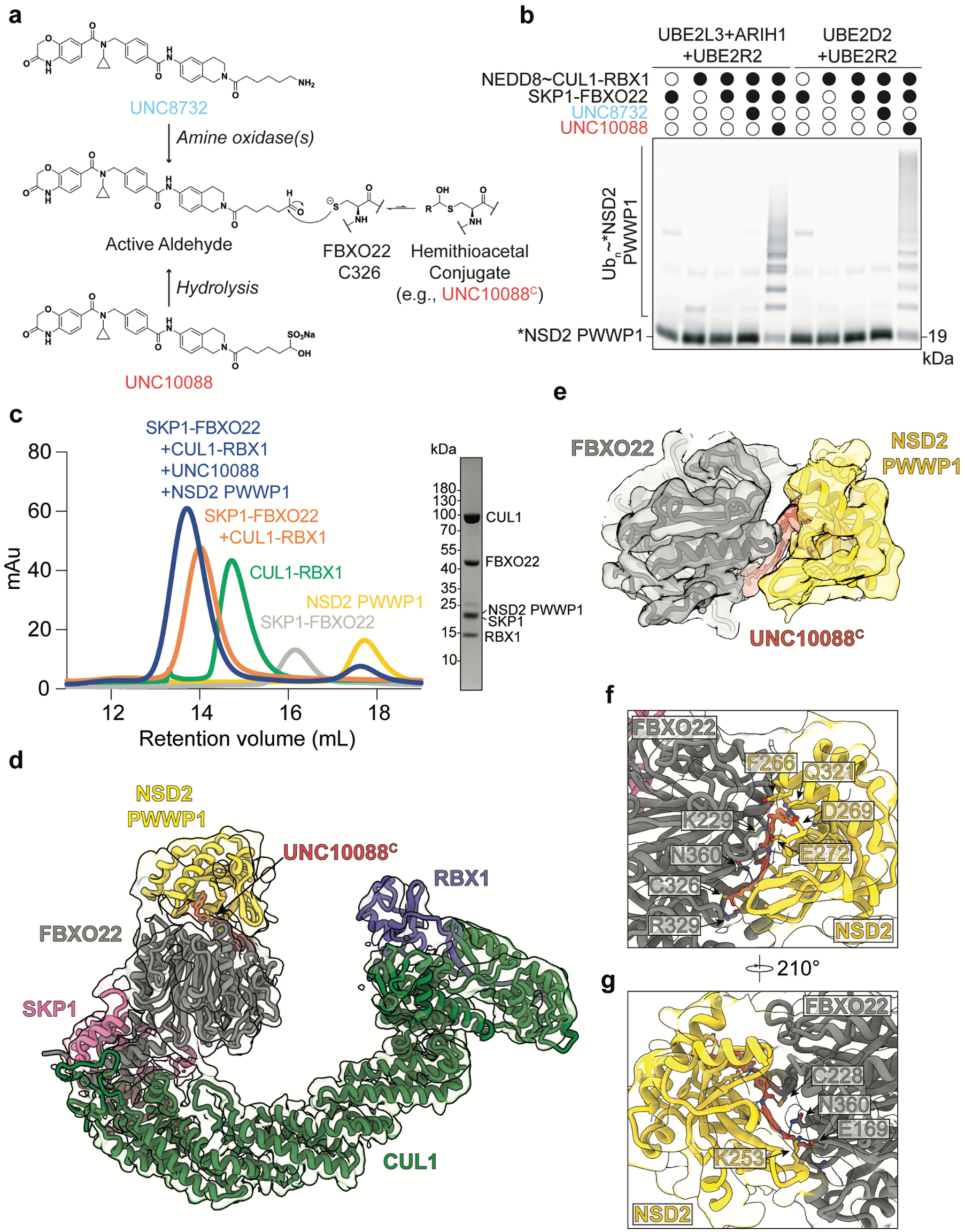
Structure of NSD2 PWWP1 bound to SCF^FBXO22^ through UNC10088. (a) Chemical structures of UNC8732 and UNC10088. UNC8732 gets metabolized by amine oxidases into a reactive aldehyde. UNC10088 contains a bisulfite warhead that gets hydrolyzed into an aldehyde under aqueous conditions that can interact with FBXO22 C326 to form a hemithioacetal conjugate (UNC10088^C^). (b) Fluorescent monitoring of an SDS-PAGE gel reveals UNC10088-mediated ubiquitination of an SCF^FBXO22^ neosubstrate *NSD2 PWWP1 using two sets of UCEs. Compounds were added at 0.5 µM. “*” indicates fluorescently labeled protein. (c) Size exclusion chromatogram (left) of the assembled SCF^FBXO22^-UNC10088^C^-NSD2 PWWP1 ternary complex for structural studies compared to the individual proteins or subcomplexes. Representative Coomassie-stained SDS-PAGE gel of the sample used for cryo-EM (right). (d) The modeled density reveals the orientation of UNC10088^C^ (orange) along the interface between FBXO22 (gray) and the neosubstrate NSD2 PWWP1 domain (yellow). SCF subunits are modeled including the adaptor protein SKP1 (pink), the inactive CUL1 scaffold protein (green), and RBX1 (blue). (e) The close-up view of the FBXO22-NSD2 PWWP1 interface reveals that UNC10088^C^ traverses a gap between the proteins. (f) A close-up view of the FBXO22-NSD2 PWWP1 interface reveals residues engaging in interactions with the adjacent protein, UNC10088^C^, or both. (g) A close-up view similar to (f) but rotated 210° about the z-axis. Additional residues on both the FBXO22 side and NSD2 side are shown to potentially engage interactions with residues of the adjacent protein, UNC10088^C^, or both.

Herein, we utilized UNC10088 to provide key fundamental insights into the repurposing of SCF^FBXO22^ for TPD through cryo-EM and detailed biochemical analysis. First, structural data with the aldehyde derivative of UNC10088 reveals a conformational change in FBXO22 that further exposes the reactive cysteine, C326. Density protruding from C326 suggests the aldehyde derivative is conjugated to C326 (henceforth annotated as UNC10088^C^), forming the previously proposed hemithioacetal linkage (Figure 1A). Additional surfaces between FBXO22-UNC10088^C^-NSD2 reveal an interaction network supporting a subtle, but significant, change in cooperative binding. Subsequent medicinal chemistry efforts reveal the ability of a benzaldehyde warhead to similarly engage C326 to promote potent NSD2 degradation, likely due to the flexibility of the Y390-containing loop near C326. Furthermore, instead of competing for native substrate recruitment, we reveal that FBXO22 can simultaneously recruit and promote the degradation of neosubstrates such as NSD2 and its native substrate BACH1. Overall, our work provides important mechanistic insights into the repurposing of FBXO22 for TPD and its potential advantages, as well as next generation NSD2 degraders.

## RESULTS

### Cryo-EM Structure of the SCF^FBXO22^-UNC10088^C^-NSD2 PWWP1 Complex

As shown in our previous study, UNC8732 and UNC10088 are converted to an aldehyde derivative by amine oxidases and hydrolysis, respectively, to enable a covalent interaction with C326 of FBXO22 (Figure 1A). Consequently, UNC10088 promotes ubiquitination of the NSD2 PWWP1 domain by SCF^FBXO22^ in the presence of different combinations of ubiquitin carrying enzymes (UCEs), E2s (UBE2D2, UBE2L3, UBE2R2) and the ARIH-type of RBR (ARIH1) (Figure 1B)^9^. In contrast, the alkylamine prodrug UNC8732 has no effect *in vitro*. However, it was unclear if the ternary complex could be stably formed for structural studies because UNC10088 has bivalent characteristics, as demonstrated by a hook effect (Extended Data Figure 1A). Therefore, we assayed complex formation between SCF^FBXO22^, UNC10088^C^, and NSD2 by size exclusion chromatography (SEC). Encouragingly, all components of the SCF^FBXO22^ -NSD2 PWWP1 complex co-migrated together and eluted off the column earlier than the individual components upon incubation with UNC10088, indicating that the complex was larger than SCF^FBXO22^ alone or any of the other individual proteins or complexes that could have formed during this process (Figure 1C).

The complex was then subjected to single particle cryo-EM analysis, allowing for 3D reconstructions at 3.8 Å resolution (Extended Data Figure 2). Based on prior structural work^12,13,19^, we were able to confidently model the SCF^FBXO22^ components and NSD2 into the cryo-EM map (Figure 1D). The entire UNC10088^C^ degrader, including the flexible alkyl chain, could also be reliably built and seems to transverse a gap at the FBXO22-NSD2 PWWP1 interface (Figure 1E). Expectedly, UNC10088^C^ interactions with NSD2 are similar to those with the NSD2 chemical probe (UNC6934) (Extended Data Figure 1B)^19^. Furthermore, additional density was observed protruding from FBXO22 Cys326, which we postulate to be a hemithioacetal linkage to the NSD2 degrader (Figure 1F). Interestingly, the NSD2 PWWP1 domain and the FBXO22 FIST domain appear to form additional interactions, potentially stabilizing the complex for potent NSD2 degradation (Figure 1F-G).

### FBXO22-NSD2 interactions facilitate cooperative binding for efficient UNC10088-dependent ubiquitination

Based on this structure, we made several mutants to test the contributions of individual interactions. First, reactive Cys326 of FBXO22 is suggested to form a covalent hemithioacetal with the UNC10088 aldehyde derivative and is therefore expected to be critical for complex formation. Based on the structure, we hypothesized that the FIST domain alone recapitulates the interaction with NSD2 in a C326-dependent manner. Performing size exclusion chromatography with the FBXO22 FIST domain and the NSD2 PWWP1 domain, indeed the complex could only be formed in a Cys326-dependent manner as a single C326A substitution ablated complex formation (Figure 2A). Additionally, the reintroduction of Cys326 into a cysteine-less version of the FIST domain (Cys326 only) rescued the defect in complex formation, further supporting that UNC10088 selectively engages Cys326 over the other 7 cysteines in the FIST domain.

**Fig. 2:**
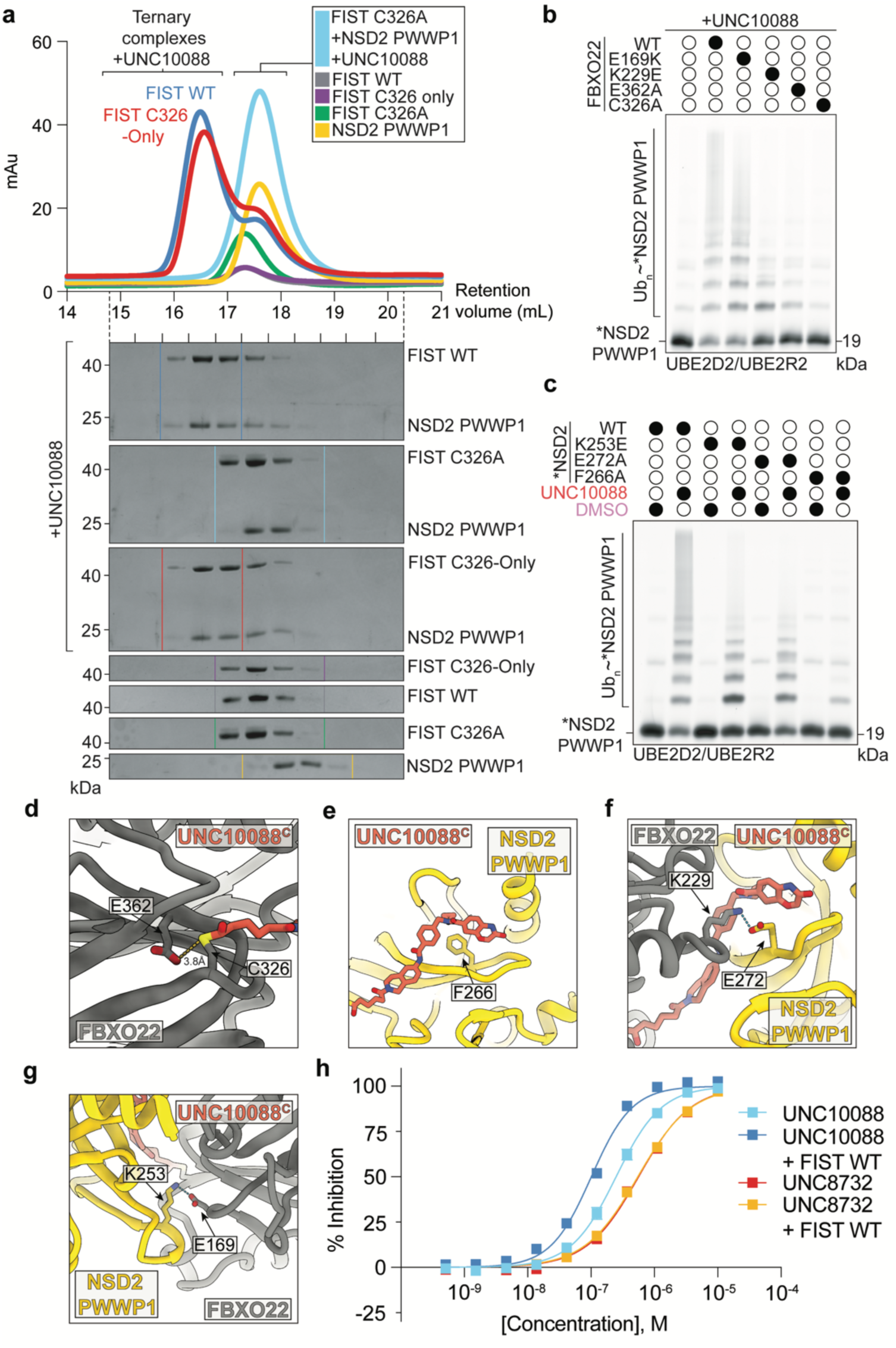
Determinants for NSD2 ubiquitination by SCF^FBXO22^ and UNC10088. (a) C326 is necessary for UNC10088-mediated FBXO22-NSD2 complex formation. Gel filtration chromatography elution profiles and corresponding Coomassie-stained SDS-PAGE gels of FBXO22 FIST wild-type or variants alone or mixed with UNC10088 and NSD2. (b) Fluorescent scan of an SDS-PAGE gel comparing the effects of indicated substitutions at FBXO22 surface residues impede UNC10088(0.5 µM)-dependent *NSD2 PWWP1 ubiquitination. (c) Similar to b. Indicated substitutions on *NSD2 PWWP1 ubiquitination by SCF^FBXO22^ and 0.5 µM UNC10088. (d-g) Close-up views of indicated residues at the interaction surface between FBXO22-UNC10088^C^-NSD2. (h) UNC10088 potently binds NSD2 PWWP1 (IC_50_ = 248 ± 22 nM) in a competitive TR-FRET assay, and interaction is enhanced in the presence of FBXO22 FIST (IC_50_ = 101 ± 8 nM), demonstrating cooperative binding (α = 3.3). A similar effect is not observed with UNC8732. *n* ≥ 3 independent experiments. Error bars: standard error of the mean.

Second, variants of either NSD2 or FBXO22 were made to explore their ability to disrupt UNC10088- and SCF^FBXO22^-dependent NSD2 ubiquitination i*n vitro* (Figure 2B-C, Extended Data Figure 1C). To begin, we focused on residues that are likely necessary for covalent conjugation or are expected to directly interact with UNC10088^C^. For example, while C326 is responsible for the nucleophilic attack of the aldehyde group of UNC10088, E362, which is adjacent to C326, may increase its activity (Figure 2D).

Consistent with this hypothesis, FBXO22 variant E362A ablated NSD2 ubiquitination (Figure 2B). On the NSD2 ligand binding interface, F266 is a core residue of the PWWP1 aromatic cage that is necessary for NSD2 ligand binding (Figure 2E, Extended Data Figure 1D). Consistently, the F266A NSD2 PWWP1 mutation also dramatically reduced SCF^FBXO22^-dependent ubiquitination of NSD2 (Figure 2C).

An exciting observation was the presence of direct interactions between NSD2 and FBXO22 that are unique to ternary complex formation. For example, an interaction network between UNC10088^C^, E272 of NSD2, and K229 of FBXO22, as well as a salt bridge between K253 of NSD2 and E169 of FBXO22, were observed in the structure (Figure 2F-G). NSD2 (E272A and K253E) and FBXO22 (K229E and E169K) variants at these surfaces exhibited reduced SCF^FBXO22^-dependent NSD2 ubiquitination (Figure 2B-C). Importantly, these defects in NSD2 ubiquitination were observed when either set of UCEs, UBE2D2/UBE2R2 (Figure 2C) or UBE2L3/ARIH1 (Extended Data Figure 1E), were used. Taken together, the mutagenesis data suggests that these additional interactions between FBXO22 and NSD2 contribute to a productive ternary complex.

Because of the direct interactions observed between FBXO22 and NSD2, we hypothesized that some degree of cooperative binding could be facilitating complex stabilization. Therefore, we developed a proximity-based Time-Resolved Fluorescence Resonance Energy Transfer (TR-FRET) assay to monitor the interaction between his-tagged NSD2 and a biotinylated NSD2 ligand, UNC7096, that is missing the modality for FBXO22-recruitment (Extended Data Figure 3A)^19^. Given the positive FRET signal observed when both NSD2 PWWP1 and UNC7096 are present, we used this setup to evaluate the displacement of UNC7096 by UNC10088 in the presence or absence of the FBXO22 FIST domain (Extended Data Figure 3A). More potent antagonism of NSD2 by UNC10088 in the presence of FBXO22 relative to UNC10088 alone would suggest a cooperative interaction between FBXO22 and NSD2. While the IC_50_ of UNC10088 alone was 248 nM, the addition of FBXO22 reduced the IC_50_ value by about 2.5-fold (IC_50_ = 101 nM), indicating positive cooperative binding with an α-factor of 3.3 (Figure 2H). In contrast, the IC_50_ value of UNC8732, which cannot bind FBXO22 *in vitro*, was unchanged in the presence of FBXO22 (∼550 nM). Importantly, this FBXO22-dependent enhancement of NSD2 antagonism by UNC10088 was absent when C326 was substituted for alanine (Extended Data Figure 3B). Taken together, the SCF^FBXO22^-NSD2 PWWP1 structure both confirms prior results and provides additional insight into the surfaces that contribute to cooperative interaction between NSD2 and FBXO22 to promote potent NSD2 polyubiquitination and degradation.

### Identification of UNC10415667, a benzaldehyde containing NSD2 degrader

While UNC10088 and UNC8732 are promising cell-free and cell-based tool compounds, respectively, we were interested in developing non-prodrug analogs that could similarly engage FBXO22 to promote NSD2 degradation without requiring bioactivation for activity. Additionally, alkyl aldehydes can be metabolically unstable and are an uncommon pharmacophore in FDA approved drugs. Surprisingly, in the previously reported FBXO22-BACH1 structure^12,13^, C326 has a relatively limited solvent accessibility due to a nearby loop containing Y390. However, our structure reveals that the loop and Y390 are shifted away to accommodate the hemithioacetal covalent intermediate between C326 and UNC10088, opening the pocket relative to the BACH1-bound FBXO22 structure (Figure 3A). Importantly, this revealed the possibility that alternative or larger warheads may be able to be accommodated to engage C326. Thus, we next designed and synthesized a set of eight compounds that contain an aromatic aldehyde in place of the aliphatic amine of UNC8732, with the expectation that these aromatic aldehydes would be more stable in a cellular environment and are easier to synthesize than their aliphatic counterparts, while potentially maintaining the ability to engage C326 in a similar reversible covalent fashion. We also hypothesized that the benzaldehyde ring may interact favorably with the side chain of nearby Y390. Within this set of compounds, we varied the length of the alkyl chain and in some cases removed it entirely (0 – 4 CH_2_), the position of the aldehyde on the aromatic ring (*ortho*-, *meta*-, or *para*-), and the size of the aromatic ring (benzene or thiophene) (Figure 3B). The compounds were tested in our *in vitro* ubiquitination assay and an NSD2 HiBiT degradation assay in U2OS cells (Figure 3C-D). Excitingly, these efforts revealed that UNC10415667 and UNC10415668 were able to induce polyubiquitination and degradation of NSD2, with UNC10415667 promoting NSD2 PWWP1 ubiquitination to the same extent as UNC10088 (Figure 3C). Similarly, treatment of NSD2 HiBit cells with 1 µM UNC10415667 for 6 hours resulted in a reduction of NSD2 levels by about 75% (Figure 3D) and did not impact cell viability (Extended Data Figure 3C). Follow up treatment of NSD2 HiBit cells with UNC10415667 in a dose response fashion revealed a DC_50_ value (concentration at which 50% of NSD2 is degraded) of 460 nM, which is about 2-fold more potent than UNC8732 and UNC10088 in the same assay (Figure 3E).

**Fig. 3:**
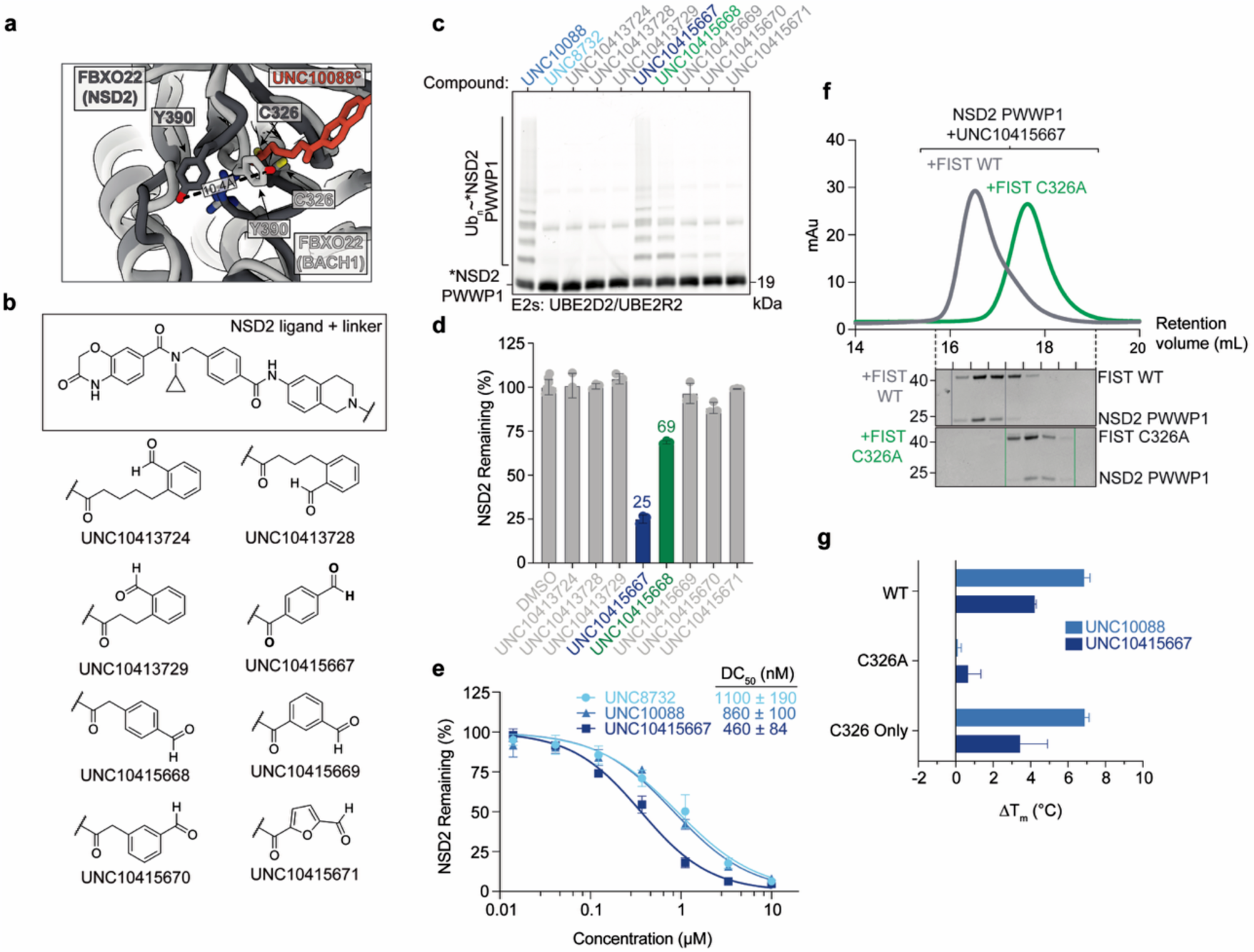
Benzaldehyde derivative UNC10415667 recapitulates SCF^FBXO22^-dependent ubiquitination and potent degradation of NSD2. (a) UNC10088^C^-induced conformational change of the loop adjacent to C326. Structural overlay of the SCF^FBXO22^-UNC10088^C^-NSD2 PWWP1 model (dark gray) and the SCF^FBXO22^-BACH1 BTB complex (light gray, PDB: 8UA3)^12^, revealing the displacement of FBXO22 residues Y390. (b) Chemical structures of UNC8732 analogs that contain an aromatic aldehyde in place of the aliphatic amine of UNC8732. (c) Benzaldehyde derivatives UNC10415667 and UNC10415668 promote FBXO22-dependent ubiquitination of *NSD2 PWWP1 when added at 0.5 µM, based on fluorescent scanning of an SDS-PAGE gel. (d) Treatment of U2OS NSD2-HiBit cells with UNC10415667 (1 µM) for 6 hours resulted in a potent reduction of NSD2 levels by about 75%. A more modest effect was observed upon treatment with UNC10415668, and all other compounds had no significant effect. *n* ≥ 3 independent experiments. Error bars: standard error of the mean. (e) Treatment of U2OS NSD2 HiBit cells with UNC10415667 in a dose response fashion revealed a DC_50_ value of 0.46 µM, which is about 2-fold more potent than UNC8732 and UNC10088. *n* = 3 independent experiments. Error bars: standard error of the mean. (f) UNC10415667 uses C326 to recruit NSD2 to the FBXO22 FIST domain as monitored by size-exclusion chromatography. Chromatograms of different elution profiles between mixtures of WT FBXO22 FIST or a C326A mutant with NSD2 and UNC10415667 (top). Representative Coomassie-stained SDS-PAGE gels of collected fractions. (g) Differential scanning fluorimetry (DSF) was used to monitor the stability of FBXO22 FIST wild type and variants in the presence of UNC10088 or UNC10415667 compared to DMSO, revealing that both compounds stabilize WT FBXO22 FIST and the C326 single cysteine variant (C326 only) as evidenced by an increase in melting temperature (ΔT_m_), but not the C326A mutant. *n* = 3 independent experiments. Error bars: standard error of the mean.

Several independent experiments revealed that the mechanism of binding and recruitment of UNC10415667 to FBXO22 is similar to UNC10088. First, in our *in vitro* ubiquitination assay UNC10415667 displayed a characteristic hook effect at similar concentrations as UNC10088 (Extended Data Figure 1B and 1F), although to a somewhat lesser degree. Second, even though there are 8 cysteines in the FIST domain of FBXO22 that could potentially react with the benzaldehyde of UNC10415667, ternary complex formation between FBXO22 FIST, NSD2 PWWP1, and UNC10415667 was dependent on the presence of Cys326 in FBXO22 (Figure 3F, Extended Data Figure 3D). Third, we used thermostability as a readout for degrader binding to FBXO22 and C326 dependence using differential scanning fluorimetry (DSF). Like UNC10088, the addition of UNC10415667 resulted in a significant increase in the melting temperature (*T*_m_) of the FBXO22 FIST domain (Figure 3G). Additionally, both compounds increased the *T*_m_ of FBXO22 in a C326-dependent manner. Specifically, when UNC10415667 was incubated with the C326A variant, a negligible shift in the *T*_m_ was observed compared to either the wild-type or the C326 only version of the FIST domain (Figure 3G). Taken together, UNC10415667 reveals that slightly larger benzaldehyde-based degraders can effectively recruit FBXO22 via C326 to promote the potent degradation of NSD2 without requiring chemical protection or bioactivation.

### Cryo-EM Structure of SCF^FBXO22^ bound to UNC10415667^C^ and NSD2

To determine if the phenyl group of UNC10415667 contributed to complex formation or promoted favorable contacts with FBXO22, we sought to determine the structure of the SCF^FBXO22^ complex bound to NSD2 through a conjugated form of UNC10415667 (UNC10415667^C^). As expected based on our size exclusion chromatography results (Figure 4A), we were able to form a stable complex of SCF^FBXO22^-UNC10415667^C^-NSD2. Using single particle cryo-EM analysis, we solved the structure to 5.7 Å resolution, revealing the configuration of the entire complex (Figure 4B, Extended Data Figure 4A-F). This structure with UNC10415667^C^ was notably very similar to the FBXO22–UNC10088^C^–NSD2 complex showing a local Cα–Cα RMSD of 1.386 Å and a global RMSD of 2.259 Å (Figure 4C). Importantly, the Y390-containing loop had again pivoted away from C326, similar to the UNC10088^C^-dependent structure, yet Y390 did not appear to be making any specific contacts with the benzaldehyde of UNC10415667. Additional density protruded from Cys326, which we could assign to the added benzene ring, confirming the formation of the covalent hemothioacetal (Figure 4D, Extended Data Figure 4G-H). As many of the interactions between FBXO22, degrader (UNC10088^C^ and UNC10415667^C^), and NSD2 are conserved in both structures, we found that UNC10415667 also promoted a cooperative interaction between FBXO22 and NSD2 (Extended Data Figure 4I), similar to UNC10088 (Figure 2H). Overall, UNC10415667 conserves many of the biochemical properties and interfaces that promote productive ternary complex formation for NSD2 ubiquitination.

**Fig. 4:**
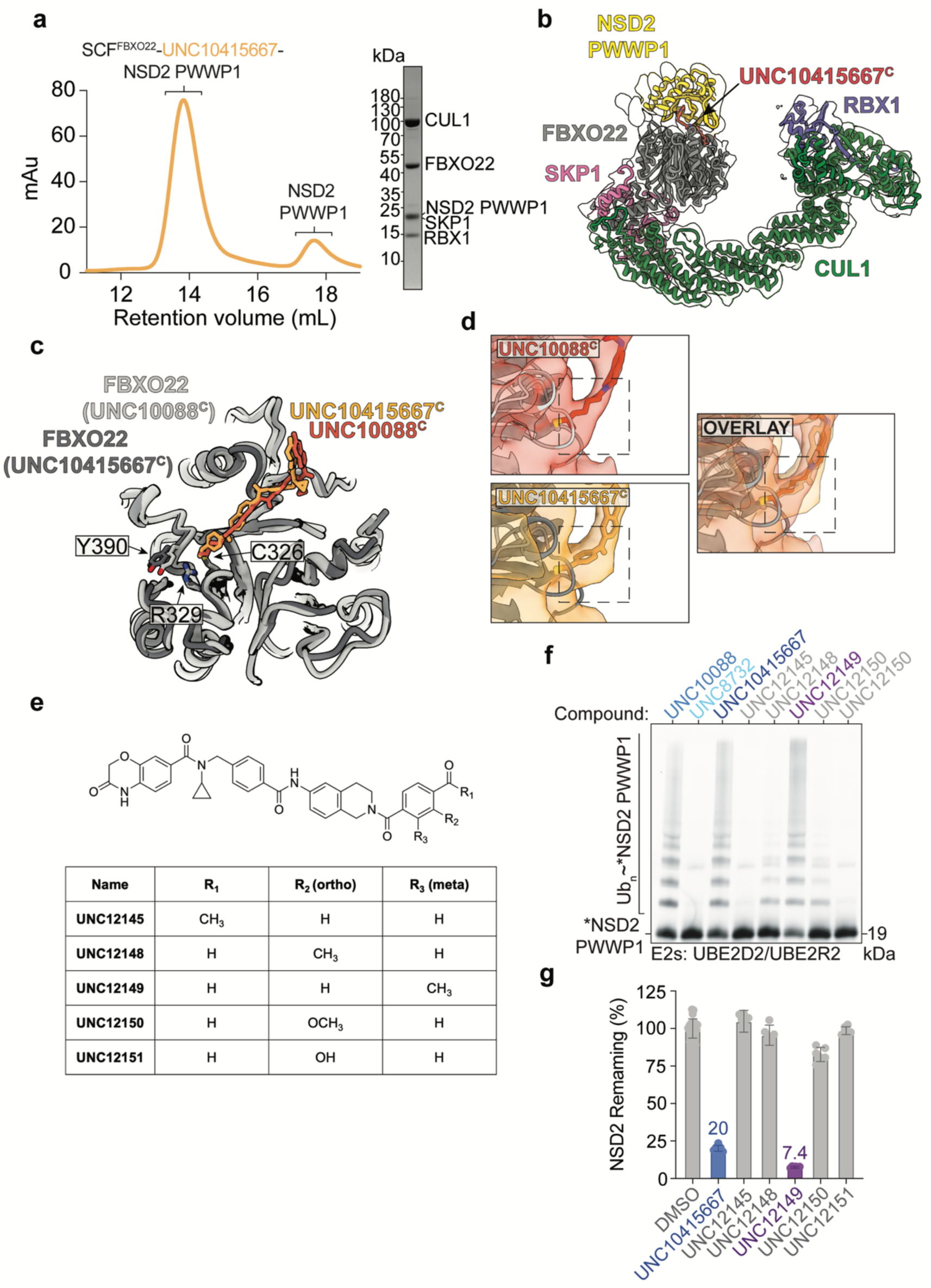
Structure of NSD2 PWWP1 bound to SCF^FBXO22^ through UNC10415667 and evaluation of UNC10415667 analogs. (a) Gel filtration profile of sample containing the assembled SCF^FBXO22^-UNC10415667-NSD2 PWWP1 complex (left) used for structural determination by cryo-EM and its representative Coomassie-stained SDS-PAGE gel (right). (b) The cryo-EM density and model reveal UNC10415667^C^ (orange) occupying a similar orientation compared to UNC10088^C^ along the interface between FBXO22 (gray) and the neosubstrate NSD2 PWWP1 domain (yellow). SCF subunits are modeled, including the adaptor protein SKP1 (pink), the inactive CUL1 scaffold protein (green), and RBX1 (blue). (c) Comparison of UNC10088^C^ and UNC10415667^C^ bound to FBXO22. (d) Additional density near C326 is observed with UNC10415667^C^ when compared to UNC10088^C^ due to the benzene ring. (e) Chemical structures of benzaldehyde compounds inspired by UNC10415667 that were tested in *in vitro* ubiquitination (f) and cell-based HiBit assays (g). (f-g) Assessment of UNC10415667 analogs by *in vitro* NSD2 ubiquitination assays, tested at 0.5 µM (f), and NSD2 degradation in our HiBit cell line (1 µM, 6 hrs) (g), revealing UNC12149 as a potent NSD2 degrader. *n* ≥ 6 independent experiments for the HiBit assays. Error bars: standard error of the mean.

### Structure-activity relationships of UNC10415667

We next designed and synthesized a few additional analogs of UNC10415667 to better understand how substitutions on the benzaldehyde ring could potentially impact FBXO22 engagement and therefore NSD2 degradation. In addition to influencing steric accessibility to the C326 binding pocket, such substituents could also potentially affect aldehyde reactivity through electronic effects. We introduced a methyl group ortho (R2; UNC12148) and meta (R3; UNC12149) to the aldehyde, as well as electron donating methoxy (UNC12150) and hydroxy (UNC12151) groups ortho to the aldehyde (Figure 4E). The ketone analog of UNC10415667 was also synthesized (UNC12145) as a negative control as the ketone is unable to covalently engage C326 in the same fashion as the corresponding aldehyde. UNC12149 which contains a methyl group meta to the aldehyde was the only analog that demonstrated significant *in vitro* NSD2 ubiquitination or NSD2 degradation in the HiBit assay, resulting in a greater than 90% reduction of NSD2 at 1 µM (Figure 4F-G, Extended Data Figure 4J). This result was further confirmed by western blot in U2OS cells, where NSD2 was undetectable upon treatment with 300 nM UNC12149 for 6 hours (Extended Data Figure 5A).

As all of the ortho substituents (R2 position) were not well tolerated, leading to little or no NSD2 degradation, we turned to our UNC10415667 structure to better understand the structure-activity relationships observed. By analyzing the cryo-EM structure, we noticed that the carbon ortho to the aldehyde is close to G359 and C326, indicating the potential for substituents at this position (e.g., methyl, hydroxy, methoxy) to introduce a steric clash or sterically hinder the interaction between the C326 side chain and the aldehyde warhead. Also, the polarity and electron donating capabilities of the methoxy and hydroxy substituents could also influence binding and aldehyde reactivity. In contrast, the methyl group at the meta(R3)-position in UNC12149 could have more space based on the structure and even potentially form van der waals interactions with nearby residues (e.g., V327, Extended Data Figure 5B). Overall, these SAR studies using the ternary complex structure including FBXO22, NSD2, and our next-generation NSD2 degrader, UNC10415667, revealed that UNC12149 is an equally potent NSD2 degrader.

### Proximity-induced degradation of NSD2 does not perturb FBXO22-dependent BACH1 degradation

A potential off-target concern of small molecule degraders is that repurposing the E3 Ub ligase will disrupt the ubiquitination and degradation of its native substrates. However, a structural comparison between the SCF^FBXO22^ structures bound to either its native substrate BACH1 or its neosubstrate NSD2 reveal that they bind completely different surfaces^12,13^ (Figure 5A). BACH1 interacts with a disordered region of FBXO22 that is facing towards CUL1 (residues 371–381; Figure 5A-B). FBXO22 C326 and NSD2 are located on an adjacent surface with no apparent overlap. These structures suggest that the recruitment of neosubstrates to SCF^FBXO22^ via Cys326 would not perturb the destruction of the native substrate BACH1.

**Fig. 5:**
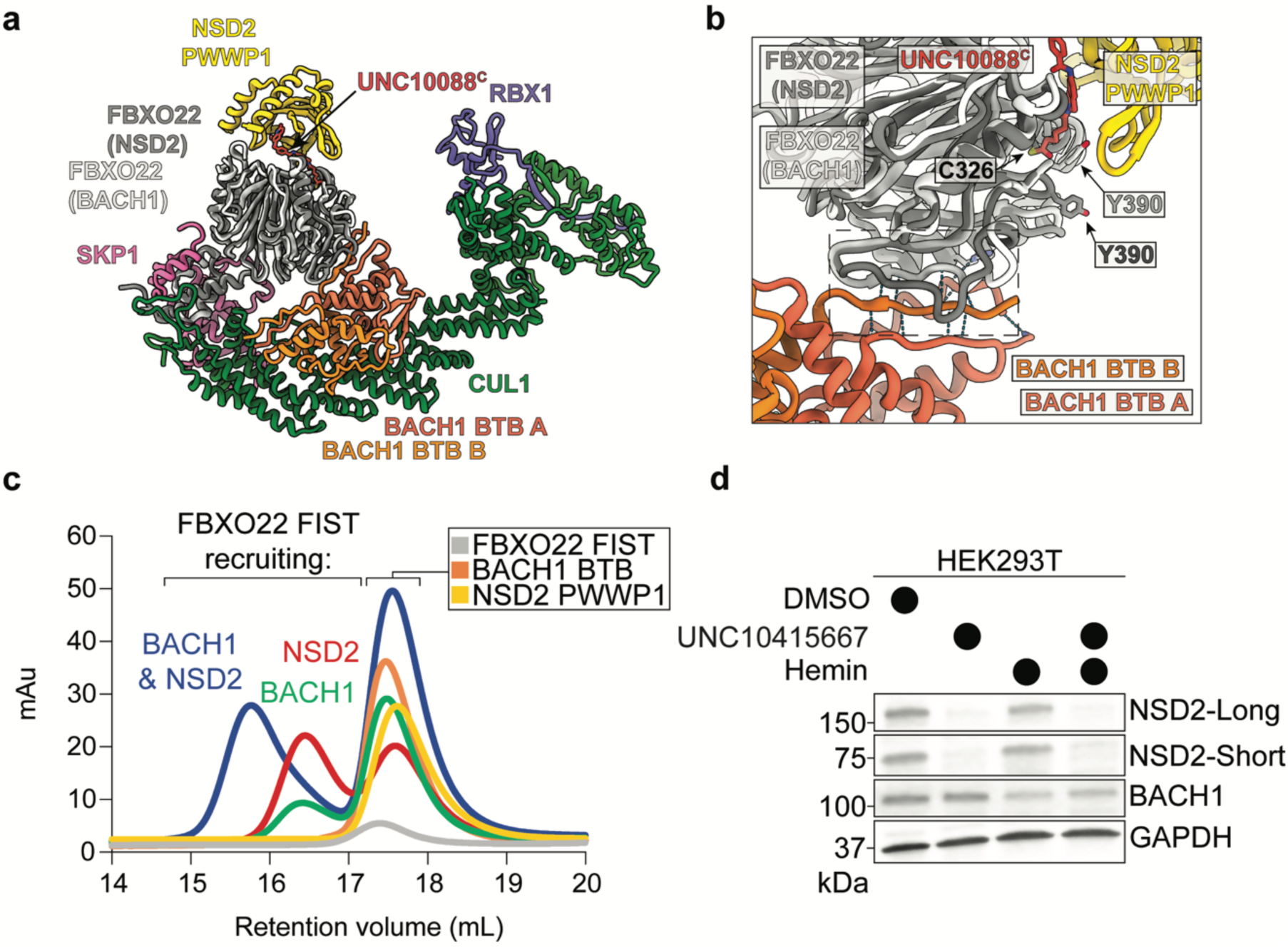
Co-recruitment of the neosubstrate NSD2 and native substrate BACH1 by SCF^FBXO22^. (a) Structural overlay of SCF^FBXO22^-UNC10088^C^-NSD2 PWWP1 and SCF^FBXO22^-BACH1 BTB (orange) (PDB: 8UA6), revealing the different binding surfaces of FBXO22^12^. (b) Close-up view of the subtle changes in the BACH1-interacting surface of FBXO22 between the two structures. (c) FBXO22 concurrently binds the neosubstrate NSD2 PWWP1 through UNC10415667 and its native substrate BACH1 BTB. Gel filtration profiles of individual proteins (FBXO22 FIST (gray), NSD2 (yellow), BACH1(orange)), subcomplexes (FBXO22 FIST-UNC10415667-NSD2 (red) or FBXO22 FIST-BACH1 BTB (green)), or FBXO22 FIST bound to UNC10415667-NSD2 and BACH1 (blue). (d) Immunoblots revealing the degradation of NSD2 and BACH1 when both 10 µM UNC10415667 and 20 µM Hemin are added to HEK293T cells for 6 hours.

To test this hypothesis, we turned to our recombinant system and purified the BACH1 BTB domain (residues 7-128) that was used in previous cryo-EM studies^12,13^. First, we performed size-exclusion chromatography on FBXO22 FIST, UNC10415667, NSD2, and BACH1. Interestingly, all of these components formed a tight complex that coeluted off the column earlier than the individual proteins, the FBXO22 FIST-BACH1 complex, or the FIST-UNC10415667-NSD2 complex (Figure 5C, Extended Data Figure 5C). To test the impact of our compounds on native and neosubstrate levels in a cell-based system, we treated HEK293T cells with UNC10415667 (10 µM for 6 hrs). As expected, both the long and short form of NSD2 were completely degraded by UNC10415667 (Figure 5D). In contrast, BACH1 levels were unchanged upon the addition of UNC10415667. As reported previously, BACH1 degradation was promoted by the addition of 20 µM Hemin^12,13,20^. When cells were treated with both UNC10415667 and Hemin, both BACH1 and NSD2 were degraded (Figure 5D). These results are consistent with the global proteomics data from our prior work demonstrating that UNC8732 specifically degrades NSD2 and does not stabilize other known FBXO22 substrates^9,21^. Although structural insights into other FBXO22 native substrates are limited, our biochemical and cell-based data further support the notion that recruiting neosubstrates to the FBXO22 surface via degraders that engage Cys326 is unlikely to significantly perturb native substrate degradation.

## DISCUSSION

Induced proximity therapeutics can provide several advantages over traditional therapeutics^2^. To this end, several ubiquitin ligases have been harnessed for targeted protein degradation (TPD), yet structural data on their ternary complexes remain scarce, limiting insight into the molecular basis of these interactions. In this example, we provide the structural basis for TPD of NSD2 by the SCF^FBXO22^ complex. As this E3 has now been harnessed to degrade several targets, elucidation of the structural basis of complex formation is essential for the optimization of existing FBXO22-recruiting degraders and in enabling the rational design of degraders for additional protein targets. A key finding of our work is that density for the postulated reversible hemithioacetal covalent bond that forms between the aldehyde active species and C326 is visible and relatively stable^9,10^. Furthermore, E362 is nearby C326 and likely keeping the cysteine reduced for covalent attack of the aldehyde as evidenced by the reduced activity upon its substitution (E362A). The structure also revealed potentially favorable contacts between NSD2 and FBXO22 upon degrader mediated complex formation, and these interactions were confirmed by mutagenesis coupled with enzymatic assays. We also demonstrated a degree of cooperative binding between the two proteins in the presence of UNC10088. While we have shown that UNC10088 and related analogs can engage both NSD2 and FBXO22 independently in a bivalent fashion, it is likely that the observed cooperative binding contributes to the potency of NSD2 degradation.

Excitingly, we also identified two novel benzaldehyde based NSD2 degraders, UNC10415667 and UNC12149, that are equally effective in cells as our previously reported prodrug degrader, UNC8732. While UNC8732 requires metabolism to reveal the active aldehyde species, such a conversion is not required with our newly identified benzaldehyde compounds; however, more detailed studies are required to fully understand if one approach is highly advantageous over the other, particularly in vivo. Importantly, these next generation compounds also suggest that the FBXO22 C326 pocket has some flexibility to accommodate groups larger than an alkyl aldehyde and that other approaches to engage this important nucleophilic residue may be possible.

A common approach to uncovering new E3 ligandable sites has been to understand where E3 ligases bind their endogenous substrates and then target those surfaces. Our discovery of a binding site on FBXO22 that is different than that of BACH1, and potentially other native targets, further emphasizes that recruiting E3s via allosteric sites is possible. Specifically, the known surfaces of FBXO22 that interact with BACH1 and BAG3, another known SCF^FBXO22^ substrate, do not overlap with the region of FBXO22 that contains C326 and engages NSD2 in our structures with either UNC10088 or UNC10415667^12,13,22^. Furthermore, our cell-based data reveals that both BACH1 and NSD2 can bind FBXO22 and be targeted by ubiquitination concurrently. This concept is consistent with the fact that TPD is a catalytic process and there are multiple different UCEs, the E2s and ARIH-type of RBRs, that cooperate with CRL ligases to create optimized geometries for ubiquitin transfer. By overlaying the structures of FBXO22-NSD2 or FBXO22-BACH1 to SCF complexes trapped mimicking ubiquitin transfer from the respective UCE (UBE2D2 or ARIH1) to substrate, we observe that NSD2 and BACH1 do not sterically clash with the UCEs (Extended Data Figure 6)^12,23-25^. Furthermore, the various orientations of both the substrates and UCEs in the modeled structures suggest that the different UCEs may have preferences to transfer ubiquitin to one type of substrate (e.g., neosubstrate) over another (e.g., native). This plasticity of the ubiquitination machinery is ideal for enabling degradation of both E3 endogenous substrates and neosubstrates in a non-mutually exclusive fashion.

Alternatively, there may be native FBXO22 substrates that are potentially recruited to the E3 ligase via engagement of Cys326 in a similar fashion to our NSD2 degraders that have not yet been uncovered. Other E3s have been shown to recruit chemically modified degrons and even non-protein substrate. For example, C-terminal amides and cyclic imides are targeted by the Cullin-RING ligases FBXO31 and CRBN, respectively^26,27^. Additionally, multiple E3s have been shown to target non-protein substrates, e.g., lipopolysaccharide^28,29^. Therefore, FBXO22 may have biological roles that have yet to be discovered where the surface containing Cys326 is needed. For example, if the chemistry of our current FBXO22 ligands is an indication of potential biological functions, FBXO22 could be playing a role in oxidative stress sensing by recognizing protein aldehydes^30,31^. FBXO22 and other ubiquitination machinery are already known to play a vital role in this pathway and the reactivity of cysteines are frequently altered^12,32,33^. Given the potential use of FBXO22-recruiting degraders as therapeutics, future studies may inform whether these degraders impact other FBXO22-dependent substrates and biological functions.

FBXO22 is overexpressed in numerous cancers based on TCGA patient data, making it a prime E3 for TPD. Additionally, NSD2 remains an attractive therapeutic target in a variety of cancer types. Our newly developed benzaldehyde-based NSD2 degraders and structural data will hopefully help guide the continued development of potent and efficacious NSD2 degraders and provide guiding principles for the design of new FBXO22 recruiting degraders for novel targets.

## ACKNOWLEDGEMENTS

We thank Shiyun Cao, Ning Zheng, and Brenda Schulman for providing constructs for SCF and UCE constructs for protein purification. The UNC Eshelman School of Pharmacy NMR Facility was supported by NIH grant S10OD032476. Our work is supported by CIHR PG-506801 (CHA); NIH R35GM153250 (MJE); Boehringer Ingelheim (DH); NIH R01CA242305 (LIJ); and NIH R35GM128855 and UCRF (NGB).

## AUTHOR CONTRIBUTIONS

TL made the NSD2-targeting molecules with advice from JLC and LIJ. KCR prepared the reagents for binding and enzyme assays. KCR, AF, and TL performed the ubiquitination assays, TR-FRET experiments, and DSF assays, respectively. KCR and IG prepared the samples for cryo-EM. KCR and SJA performed the structural analysis with advice from DH and NGB. TL and XW performed the cell-based degradation assays with advice from LIJ and MJE. CHA, JLC, MJE, LIJ, and NGB conceived the project. KCR, TL, SJA, DH, LIJ, and NGB wrote the manuscript with assistance from all authors.

## DATA AVAILABILITY

EM density maps and all raw data will be deposited to the EMDB. The corresponding atomic coordinates will be deposited in the RCSB Protein Data Bank. Additional data available upon request.

## EXTENDED DATA FIGURES and FIGURE LEGENDS

**Extended Data Fig.1:**
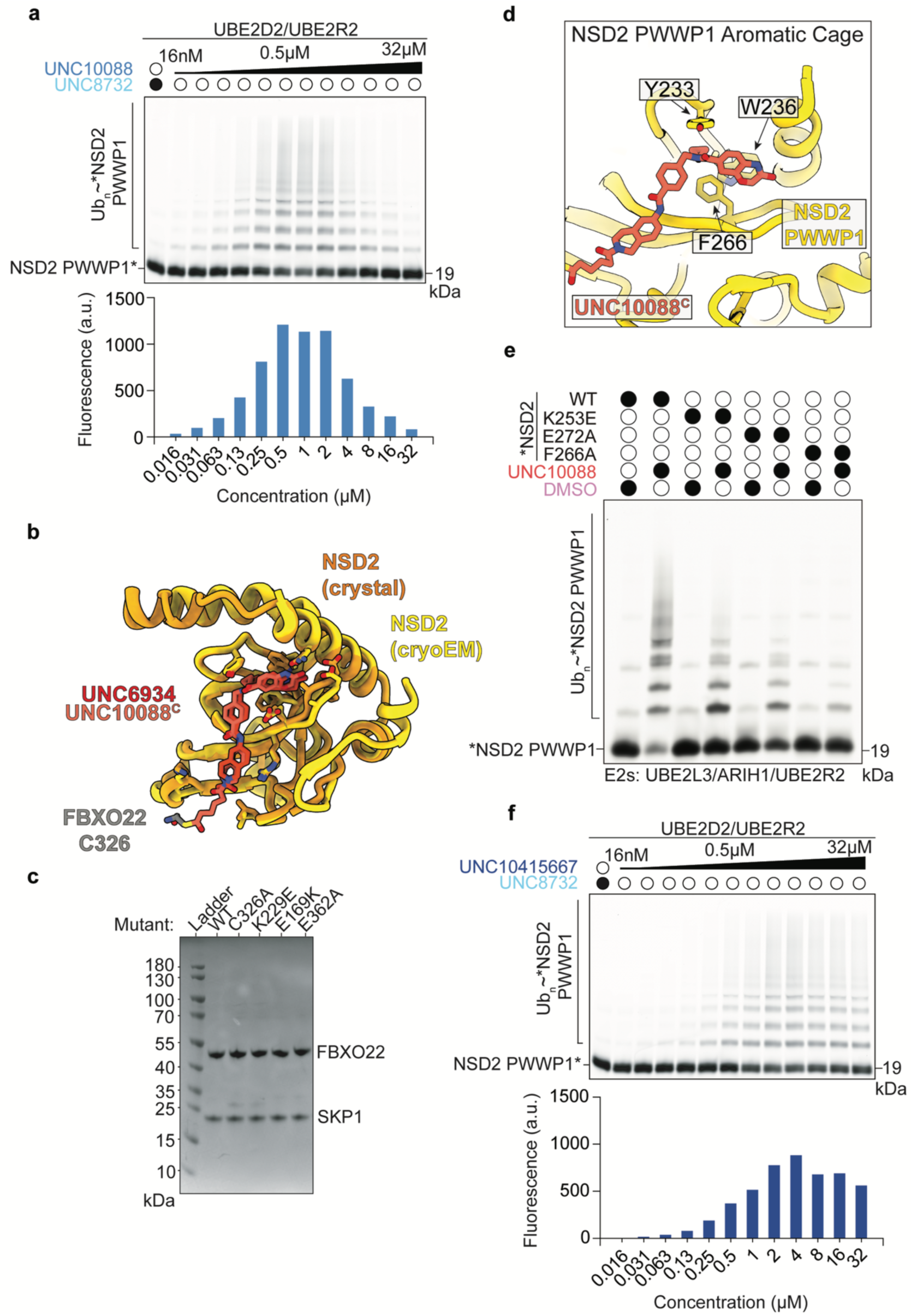
Structural comparisons using the SCF^FBXO22^-UNC10088^C^-NSD2 PWWP1 cryo-EM model reveal UNC10088^C^ occupies a similar orientation relative to previous structures and representative gels related to ubiquitination assays in Fig 2. (a) Fluorescent monitoring of an SDS-PAGE gel (top) and quantified fluorescence from ubiquitination of NSD2 (bottom) demonstrating an inhibitory hook effect upon titration of UNC10088 in SCF^FBXO22^-degrader-dependent ubiquitination. Fluorescence readout was quantified by performing a background subtraction of the negative control fluorescence (0.5 µM of UNC8732) from each condition. (b) Structural comparison of SCF^FBXO22^-UNC10088^C^-NSD2 PWWP1 complex and the prior NSD2 crystal structure bound to the ligand used to derive UNC10088 (UNC6934) and similar molecules for FBXO22-mediated degradation^19^. (c) SDS-PAGE gel of SKP1-FBXO22 WT and variants tested in Figure 2B. normalized to 5 µM. Proteins bands were visualized using Coomassie stain. (d) Aromatic cage of NSD2 for ligand recruitment. (e) Similar to Fig. 2C, fluorescent monitoring of an SDS-PAGE gel comparing the effects of indicated substitutions on *NSD2 PWWP1 ubiquitination by SCF^FBXO22^ and UNC10088 using UBE2L3, ARIH1, and UBE2R2 as the UCEs. (f) Fluorescent monitoring of an SDS-PAGE gel (top) and quantified fluorescence from ubiquitination of NSD2 (bottom) demonstrating an inhibitory hook effect upon titration of UNC10415667 in SCF^FBXO22^-degrader-dependent ubiquitination. Fluorescence readout is quantified by performing a background subtraction of the negative control fluorescence (0.5 µM of UNC8732) from each condition.

**Extended Data Fig. 2:**
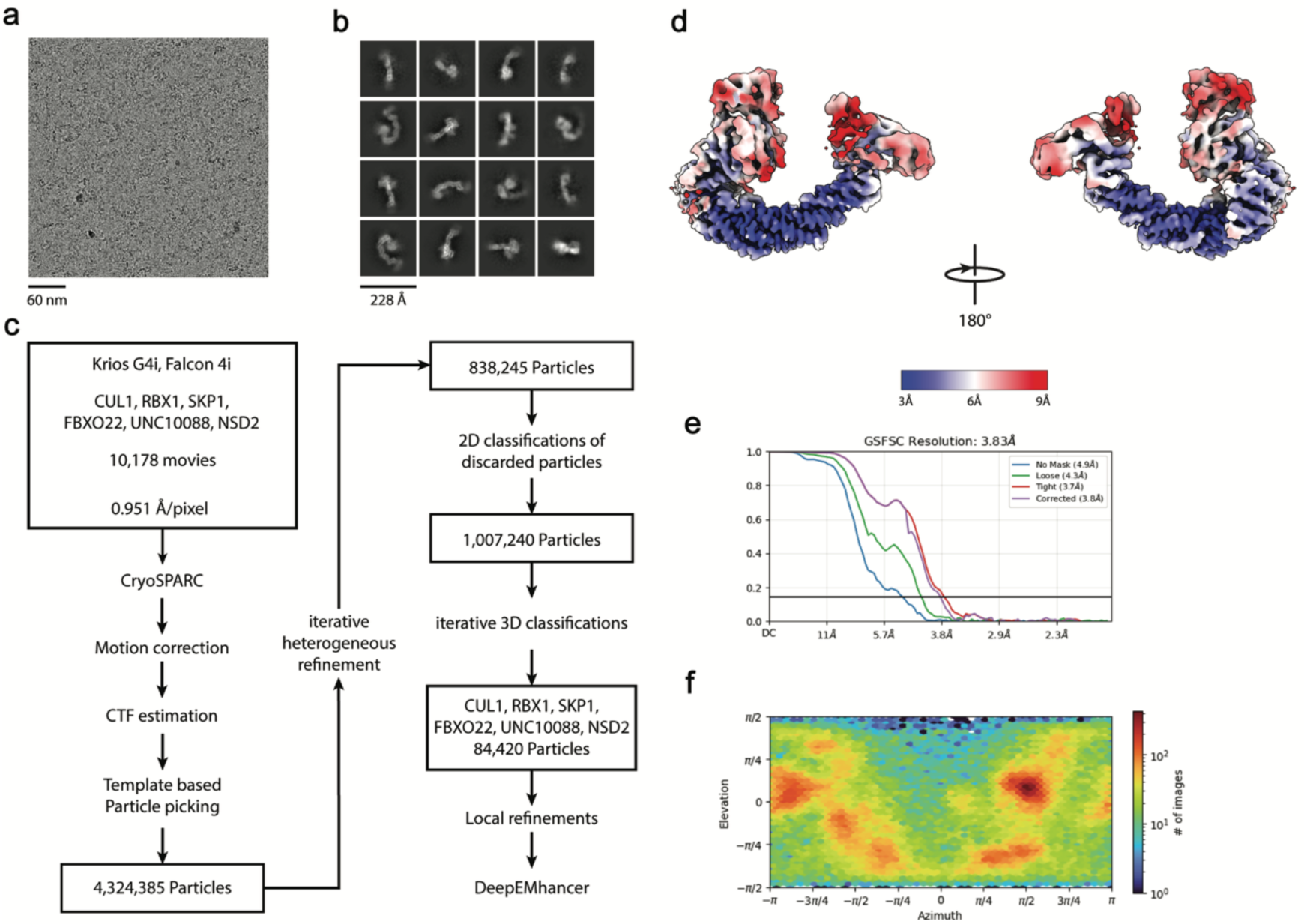
Cryo-EM analysis of the SCF^FBXO22^-UNC10088^C^-NSD2 PWWP1 complex. (a) Representative micrograph. (b) Representative 2D class averages. (c) Processing pipeline. (d) Local resolution estimation of local refined map. (e) GSFSC curve of local refined map. (f) Viewing angle distribution.

**Extended Data Fig. 3:**
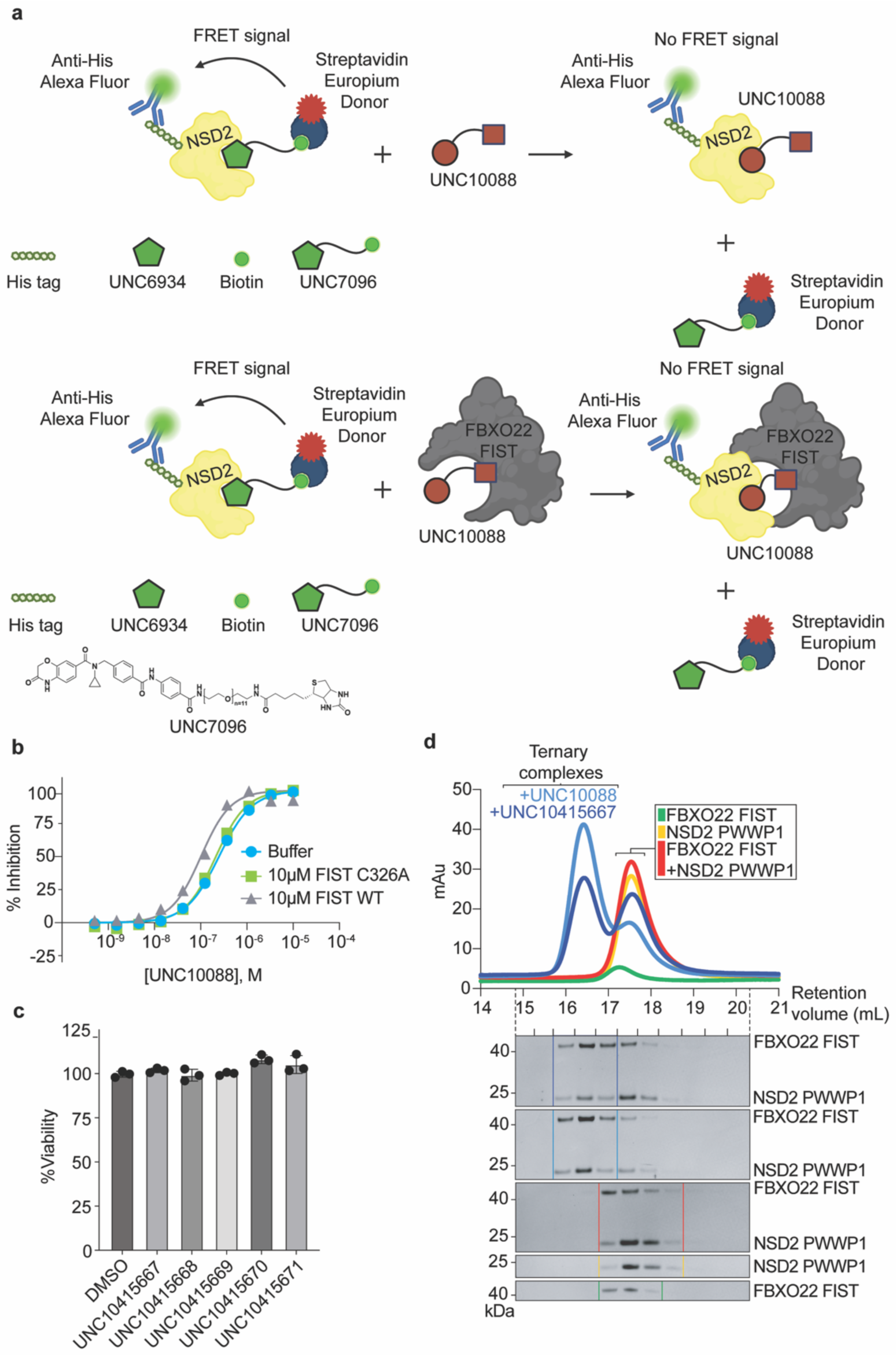
Cooperative inhibition of UNC10415667 and FBXO22 using TR-FRET. (a) Cartoon model of TR-FRET competition assay monitoring the interaction between His-tagged NSD2 PWWP1 and a biotinylated version of the NSD2 PWWP1 ligand (UNC7096), which is disrupted by the addition of UNC10088 in the presence or absence of FBXO22 FIST. (b) Competitive inhibition of the TR-FRET signal upon addition of UNC10088 in the presence of buffer (IC_50_ = 248 ± 22 nM), WT FIST (IC_50_ = 101 ± 8 nM), or C326A FIST (IC_50_ = 227 ± 10 nM). Curve fits of UNC10088 titrations with or without the FBXO22 FIST domain showing cooperative binding are repeated from Fig 2H for comparison with the FIST domain harboring the C326A variant. *n* ≥ 3 independent experiments. Error bars: standard error of the mean. (c) Representative bar graph of percent (%) cell viability of U2OS cells treated with benzaldehyde derivatives of UNC8732 from Fig 3B. *n* = 3 independent experiments. Error bars: standard error of the mean. (d) UNC10088- and UNC10415667-mediated FBXO22 FIST-NSD2 PWWP1 ternary complex formation, similar to Fig. 2A. Gel filtration chromatography elution profiles of FBXO22 FIST wild-type alone or mixed with NSD2 and either UNC10088 or UNC10415667.

**Extended Data Fig. 4:**
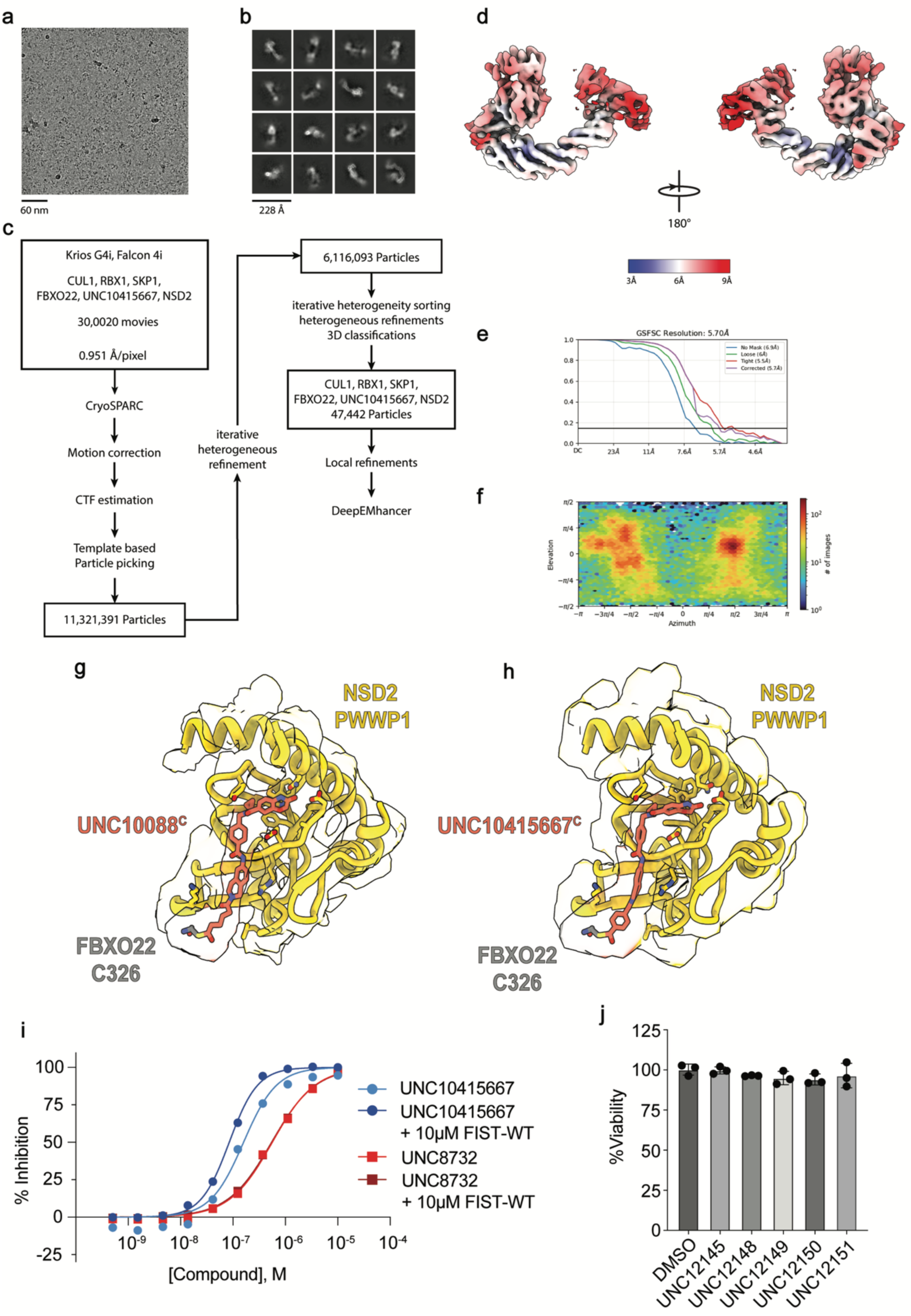
Cryo-EM analysis of the SCF^FBXO22^-UNC10415667^C^-NSD2 PWWP1 complex and additional assays with UNC10415667 and its derivatives. (a) Representative micrograph. (b) Representative 2D class averages. (c) Processing pipeline. (d) Local resolution estimation of local refined map. (e) GSFSC curve of local refined map. (f) Viewing angle distribution. (g) Close-up view of UNC10088^C^ (orange) recruitment of NSD2 PWWP1 (yellow) to FBXO22 via FBXO22 C326 (gray). (h) Close-up view of UNC10415667^C^ (orange) recruitment of NSD2 PWWP1 (yellow) to FBXO22 via FBXO22 C326 (gray). (i) Competitive inhibition of the TR-FRET signal upon addition of UNC10415667 (IC_50_ = 159 ± 24 nM), UNC10415667 in the presence of FBXO22 FIST (IC_50_ = 85 ± 4 nM), or UNC8732 in the presence or absence of FBXO22 FIST (IC_50_ ∼ 550 nM), similar to Fig. 2H. Curve fits of UNC8732 titrations with or without the FBXO22 FIST domain are repeated from Fig 2H for comparison. *n* ≥ 3 independent experiments. Error bars: standard error of the mean. (j) Representative bar graph of percent (%) cell viability of U2OS cells treated with benzaldehyde derivatives inspired by UNC10415667. *n* = 3 independent experiments. Error bars: standard error of the mean.

**Extended Data Fig. 5:**
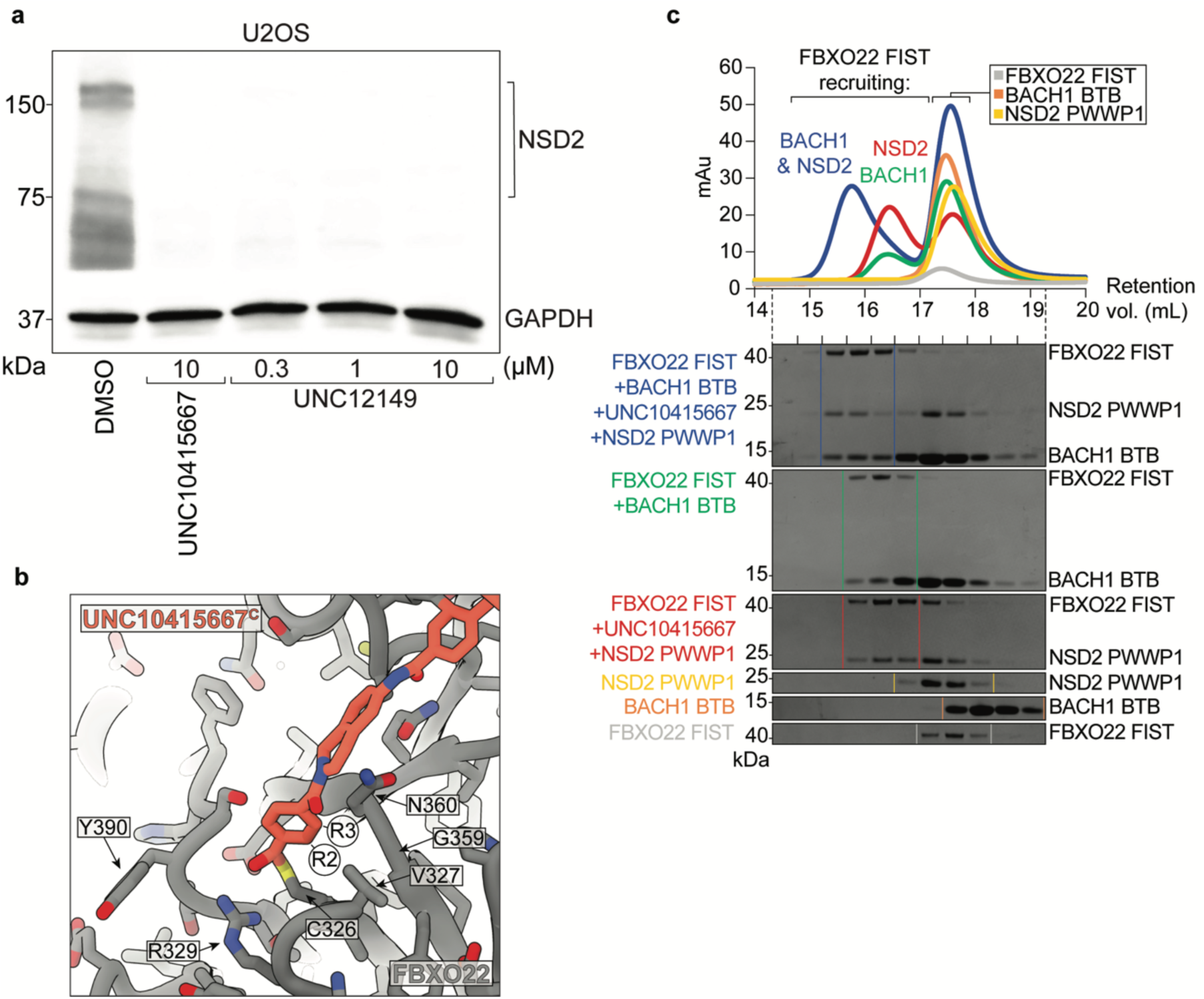
Validation of UNC10415667-, UNC12149-mediated NSD2 degradation in cells and representative gels related to filtration profiles in **Fig 5.** (a) Immunoblot of NSD2 degradation in U2OS cells after 6 hr treatment with UNC10415667 or UNC12149 versus DMSO. GAPDH is used as a loading control. (b) Close-up view of SCF^FBXO22^-UNC10415667-NSD2 structure highlighting the R2 and R3 positions (Figure 4E) and neighboring FBXO22 residues. (c) Gel filtration profiles (top) of individual proteins (FBXO22 FIST (gray), NSD2 (yellow), BACH1(orange)), subcomplexes (FBXO22 FIST-UNC10415667-NSD2 (red) or FBXO22 FIST-BACH1 BTB (green)), or FBXO22 FIST-UNC10415667-NSD2-BACH1 (blue). Representative Coomassie-stained SDS-PAGE gels of sizing fractions (bottom). Filtration profiles from Figure 5C are repeated here for clarity.

**Extended Data Fig. 6:**
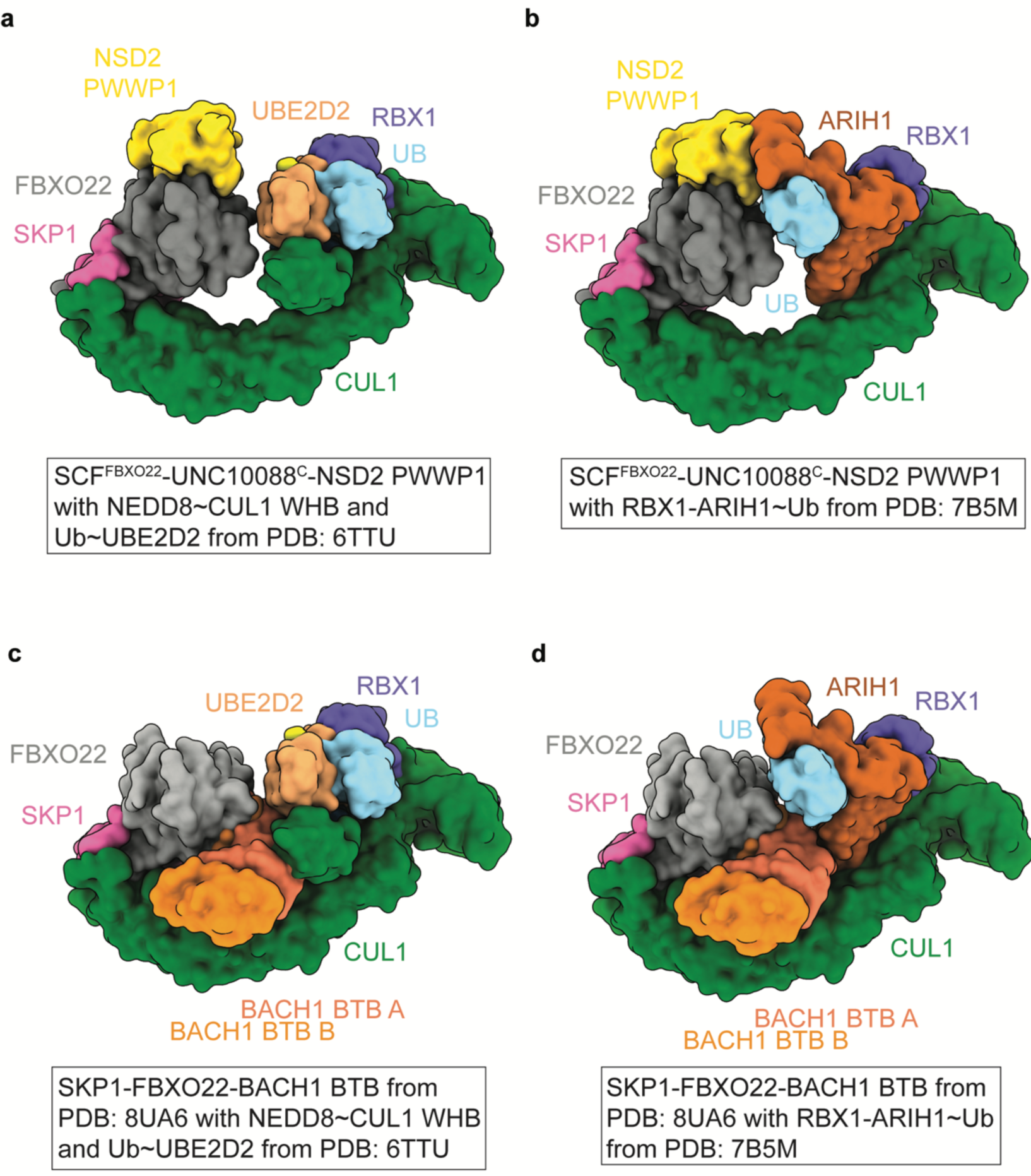
Structural models of aligning SCF^FBXO22^-UNC10088-NSD2 (A-B) or SCF^FBXO22^-BACH1 BTB (C-D) structures with active SCF structures using either UBE2D2 (A,C) or ARIH1 (B,D) as the UCE. (a) Overlay of the SCF^FBXO22^-UNC10088^C^-NSD2 cryo-EM model surface with NEDD8∼CUL1 WHB and Ub∼UBE2D2 from an active NEDD8∼CUL1-RBX1 N89R-SKP1-monomeric-βTRCPΔD-IκBɑ-Ub∼UBE2D2 complex (PDB: 6TTU)^24^. (b) Overlay of the SCF^FBXO22^-UNC10088^C^-NSD2 cryo-EM model surface with RBX1-ARIH1∼Ub from an active CUL1-RBX1-SKP2-CKSHS1-p27∼Ub∼ARIH1 complex (PDB: 7B5M)^25^. (c) Overlay of the FBXO22-BACH1 BTB cryo-EM model surface (PDB:8UA6) with NEDD8∼CUL1 WHB and Ub∼UBE2D2 from an active NEDD8∼CUL1-RBX1 N89R-SKP1-monomeric-βTRCPΔD-IκBɑ-Ub∼UBE2D2 complex (PDB: 6TTU)^24^. (d) Overlay of the FBXO22-BACH1 BTB cryo-EM model surface (PDB:8UA6) with RBX1-ARIH1∼Ub from an active CUL1-RBX1-SKP2-CKSHS1-p27∼Ub∼ARIH1 complex (PDB: 7B5M)^25^.

## MATERIAL AND METHODS

### Protein Purification

All proteins used in this study are of human origin.

Full-length 6xHis-tagged SKP1-GST-tagged FBXO22 (WT and variants) and CUL1-GST-tagged RBX1 were co-expressed in Trichoplusia ni (Tni) cells. The FBXO22 FIST constructs (WT and variants) harbored a C-terminal Twin-Strep tag and expressed without SKP1 in Tni cells. Baculovirus-infected insect cells for protein expression were cultured in ESF921 medium supplemented with 1 μg/mL amphotericin B and 5 μg/mL penicillin-streptomycin. Cells were harvested, lysed by sonication, and lysate was clarified by centrifugation. Proteins were isolated using affinity chromatography according to their respective tags, glutathione agarose 4B resin or with strep-tactin sepharose resin. After the tags were removed, via proteolytic cleavage, the SKP1-FBXO22 and CUL1-RBX1 complexes were further purified using anion exchange chromatography and cation exchange chromatography, respectively, and then by size exclusion chromatography into 20 mM HEPES pH 8, 200 mM NaCl, 1 mM DTT. The FBXO22 FIST domains were purified into the same buffer except with 0.5 mM TCEP rather than DTT.

The Split ‘n’ Co-express version of full-length CUL1-RBX1 was expressed and purified similar to prior studies^34^. Split ‘n’ co-expressed CUL1-RBX1 was neddylated as described below and used exclusively for biochemical assays while the full length CUL1-RBX1 expressed from insect cells was used for structural studies.

ARIH1, UBE2R2, NEDD8, and BACH1 BTB were expressed similarly to prior studies ^12,13,35^ with an N-terminal GST tag in BL21(DE3)-CodonPlus RIL cells in LB medium supplemented with 150 µg/mL ampicillin and 30 µg/mL chloramphenicol. GST-tagged APPBP1-UBA3 and UBC12 were expressed in BL21-Gold(DE3) cells in LB medium supplemented with 150 µg/mL ampicillin. After the sonication and centrifugations, GST-tagged proteins were isolated using affinity chromatography with GS4B resin and the GST tag was removed by TEV protease. To further purify NEDD8, UBE2R2, and BACH1-BTB, these were passed over GS4B resin to remove the liberated GST tag prior to SEC. APPBP1-UBA3 and ARIH1 were subjected to anion-exchange chromatography and UBC12 was subjected to cation-exchange chromatography prior to SEC.

NSD2 PWWP1 (WT and mutants) were expressed with an N-terminal 6xHis tag in BL21(DE3)-CodonPlus RIL cells in LB medium supplemented with 50 µg/mL kanamycin and 30 µg/mL chloramphenicol. His-tagged UBE2L3 and UBA1 were expressed similarly except with LB medium supplemented with 150 µg/mL ampicillin and 30 µg/mL chloramphenicol. After lysis and centrifugation, the proteins were isolated by using affinity chromatography with Ni^2+^-NTA agarose resin and the 6xHis tag was cleaved using TEV protease except for UBA1. To further purify NSD2 PWWP1, protein was passed over Ni^2+^-NTA resin to remove the liberated His tag. UBA1 was further purified using anion exchange chromatography. Lastly, NSD2 PWWP1 (WT and mutants), UBA1, and UBE2L3 were purified by SEC.

Un-tagged UBE2D2 and Ub were expressed in BL21(DE3)-CodonPlus RIL cells in LB medium similarly to previous studies^9^. After sonication, the clarified lysate for Ub was acidified using acetic acid to a pH of ∼4.5. Precipitation was removed by centrifugation and lysate was dialyzed overnight in 25 mM sodium acetate pH 4.5 at room temperature. UBE2D2 and Ub were isolated using cation-exchange chromatography and eluted with either 50 mM HEPES pH 7, 250 mM NaCl, 5 mM DTT or 25 mM sodium acetate pH 4.5, 250 mM NaCl, respectively. They were then subjected to SEC. The sizing buffer was 20 mM HEPES pH 8, 200 mM NaCl, 1 mM DTT unless otherwise specified, and the purified protein was flash frozen for storage at -80°C.

### SCF^FBXO22^-Degrader-NSD2 PWWP1 ternary complex formation for cryo-EM

Formation of the SCF^FBXO22^-UNC10088-NSD2 PWWP1 and SCF^FBXO22^-UNC10415667-NSD2 PWWP1 ternary complexes was performed by incubating SKP1-FBXO22, CUL1-RBX1, NSD2 PWWP1 and UNC10088 or UNC10415667 at a 1:1:1:1 molar ratio for 4 hours on ice. The incubated sample was loaded onto a Superdex 200 Increase SD200 column (Cytiva) for size-exclusion chromatography and fractions containing all subunits of the SCF^FBXO22^-Degrader-NSD2 PWWP1 complex were pooled and concentrated. The SCF^FBXO22^-UNC10088-NSD2 PWWP1 and SCF^FBXO22^-UNC10415667-NSD2 PWWP1 complexes were concentrated to 33 µM and 10.9 µM, respectively, and flash frozen.

### Size-exclusion chromatography to monitor ternary complex formation

FBXO22 WT and variants were incubated on ice with NSD2 PWWP1 and either UNC10088 or UNC10415667 at 5 µM. After 1 hour, the samples were centrifuged at 16,060xg for 10 minutes at 4°C. Each sample was examined for C326- and compound-dependent complex formation by comparing the SEC chromatograms and Coomassie-stained SDS-PAGE gels. Similar experiments were also performed in the presence of BACH1 BTB to demonstrate that both native and neosubstrates can bind simultaneously. These experiments were performed using a Superdex 200 Increase SD200 column (Cytiva) and 20 mM HEPES pH 8, 200 mM NaCl, 1 mM DTT.

### Generation of NEDD8-conjugated CUL1-RBX1

CUL1 neddylation was performed by incubating 8 µM CUL1-RBX1, 500 nM APPBP1-UBA3, 1 µM UBC12, 10 µM NEDD8, and neddylation buffer (50 mM Tris pH 7.6, 100 mM NaCl, 2.5 mM MgCl_2_, 150 uM ATP) at room temperature for 10 minutes. The reaction was quenched with 10 mM DTT and subjected to SEC.

### Fluorescent labeling of NSD2 using sortase-mediated transpeptidation

A 10X sortase buffer was prepared with 500 mM Tris pH 7.6, 1.5 M NaCl and 150 mM CaCl_2_. 50 µM of NSD2, 1 mM of the fluorescent peptide, 1 µM sortase, 1X sortase buffer were incubated on ice. Samples were quenched with 10 mM EDTA pH 8 and buffer exchanged into 20 mM HEPES pH 8, 200 mM NaCl, 1 mM DTT to remove excess fluorescent peptide, passed over Ni-NTA resin to remove excess sortase, and subjected to SEC. For NSD2 PWWP1 and its variants, a fluorescent peptide containing 5-FAM-LPETGG was used.

### Ubiquitination assays using purified components

Ubiquitination assays were performed by mixing 2 separate solutions. First, SKP1-FBXO22, fluorescently labeled NSD2-PWWP1 (*NSD2-PWWP1), and compounds, unless otherwise indicated, were pre-incubated for 15 minutes on ice. Second, UBA1, UBE2R2, UBE2D2 or UBE2L3 and ARIH1, NEDD8∼CUL1-RBX1, MgATP, and reaction buffer (20 mM HEPES pH 8, 200 mM NaCl, 0.5 mg/mL bovine serum albumin (BSA)) were combined on ice to generate a master mix. A standard volume of master mix was added to each reaction tube and Ub was added to initiate the ubiquitination reaction containing 100 µM Ub, 1 µM SKP1-FBXO22, 0.5 µM *NSD2 PWWP1, 0.5 µM compound, 0.1 µM UBA1, 1 µM UBE2R2, 1 µM UBE2D2 or 1 µM UBE2L3 and 1 µM ARIH1, 1 µM NEDD8∼CUL1-RBX1 and 5 mM MgATP in reaction buffer. Ubiquitination assays comparing *NSD2 PWWP1 mutants were performed similarly except SKP1-FBXO22, *NSD2 PWWP1 mutants, and compounds were not incubated separately prior to the start of the reaction. The reactions proceeded at room temperature for 15 minutes until it was quenched by adding SDS Loading Buffer. The ubiquitinated *NSD2 PWWP1 was visualized by subjecting the reactions to SDS-PAGE and imaging using the Amersham Typhoon.

### CryoEM screening and data collection

CryoEM grids were screened on a 200 kV Glacios TEM (Thermo Fisher) at the VBCF EM facility. Grids with good ice quality and monodispersed particles were selected for data collection on a 300 kV Titan Krios G4 TEM (Thermo Fisher) at the IMP. This instrument was equipped with an E-CFEG electron source, a Selectris X imaging filter (Thermo Fisher) with a 10eV slit width, and a Falcon 4i direct electron detector. Data collection was performed using EPU (Thermo Fisher) with 40 frames per exposure, a pixel size of 0.951 Å/px, a total dose of 40 e⁻/Å², and a defocus range of −1.0 to −2 µm in −0.2 µm increments.

### CryoEM image processing

All datasets were processed using CryoSPARC v4.6. Pre-processing included Patch Motion Correction, Patch CTF Estimation, Curate Exposures, Particle Template Picking, Particle Extraction, 2D Class Cleaning and, Ab-initio Reconstruction. Heterogeneity sorting was performed using iterative heterogeneous refinements with a good and a bad class to remove trash particles. Furthermore, local 3D classifications with a smooth mask covering FBXO22 and NSD2 was used to identify a homogeneous NSD2 bound particle subset. Final 3D refinements were done using Non-Uniform Refinements and Local Refinements with smooth masks for FBXO22 and NSD2, the N-terminal part of CUL1 and SKP1, and the C-terminal part of CUL1 and RBX1. The local refined maps were aligned to the Non-Uniform refined map, reconstructed using the vop max command in Chimerax and the composite map was post-processed using deepemhancer.

### Protein biotinylation for TR-FRET assays

20 µM FBXO22 FIST containing a C-terminal-Avitag, 10 µM BirA ligase, 40 µM biotin, and 5 mM MgATP were incubated for 1 hour on ice. Biotinylated proteins were further purified using cation-exchange chromatography to remove the BirA, followed by SEC.

### TR-FRET Assay

The TR-FRET assay was adapted to assess cooperativity from the following the workflows previously reported^36,37^. Assays were performed using white, low-volume, flat-bottom, nonbinding 384-well microplates (Greiner, Catalog No. 784904) containing a final assay volume of 10 μL per well. The assay buffer contained 20 mM Tris–HCl (pH 7.5,) and 150 mM NaCl, 0.05% Tween-20 (v/v), 2 mM DTT, and 1% DMSO (v/v).

LANCE Europium (Eu)-W8044 Streptavidin conjugate (2 nM) and LANCE Ultra ULight-anti6x-His labeled antibodies (10 nM) were used as donor and acceptor fluorophores associated with the tracer ligand and protein, respectively. Final assay concentrations of 63 nM His-tagged NSD2 PWWP1 and 31 nM of UNC7096 (a biotinylated version of the NSD2 ligand) as a tracer ligand were used for compound testing. UNC8732, UNC10088, and UNC10415667 were diluted in a 10-point, 3-fold serial dilution and indicated compounds were pre-incubated with FBXO22 FIST for 30 min. A master mix of NSD2 PWWP1, UNC7096, and TR-FRET reagents was made prior to its addition to the compounds and/or FBXO22 FIST. The plates were sealed with metallic covers, mixed gently for 1 min, and allowed to incubate in a dark space for 1 hour. After 1 hour, the plate was read on an EnVision 2103 plate reader with a 320 nm excitation filter and 615 nm/ 665 nm emission filters simultaneously with a dual mirror D400/D630 and a 100 μs delay. The raw TR-FRET signal was displayed as a ratio of acceptor/donor (665/615 nm) emission counts. Raw TR-FRET signals were normalized to a high inhibition control (10 μM of UNC10088) and negative control (1% DMSO). The normalized TR-FRET signals were plotted against the compound concentrations. These data were fitted with a four-parameter nonlinear regression analysis using GraphPad Prism software (v 10.4.1) to calculate IC50s. Ki values were calculated from the IC50s using the tight-binding model^38,39^. The cooperativity factor (α) was calculated as the ratio of Ki (Binary)/ Ki (Ternary), with α>1 indicating positive cooperativity between NSD2 PWWP1 and FBXO22 FIST based on its disruption of NSD2 binding to UNC7096.

### NSD2 HiBiT degradation assay

The NSD2 genomic sequence was obtained from the UCSC genome browser and used to design sgRNAs to introduce HiBiT tags at either the 5’ or 3’ ends of the gene. sgRNA were assembled into RNPs using TrueCut Cas9 Protein v2 (Invitrogen, Cat#A36497), an sgRNA crRNA:tracrRNA duplex (below; IDT) and ssODN (below; IDT). RNPs were introduced into U2OS cells by electroporation using a Neon XT electroporation system (1750V,10ms, 3 pulses) in R-buffer, using Thermo Fisher Neon transfection kit (Cat# MPK1025K) and according to the manufacturers protocol. After plating for recovery for four days, single cell clones limited dilution cloning. Single cell clones that grew out were identified as containing integrated HiBiT tags on NSD2 by performing HiBiT assays using HiBiT lytic assay (Promega: N3040) and read on a plate reader. Clones were further confirmed by PCR analysis.

sgRNA crRNA:tracrRNA sequence:AGAGGGCAAATAGcgccagg

ssODN sequence:

GGGAAGCCGAAGGGGAAGAGGCGGCGGCGGAGGGGCTGGCGGAGAGTCACAGAGGGCAAAAACAGGATCAGGGGCAGCAGCGGCGGCAGCAGCGGCGTGAGCGGCTGGCGGCTGTTCAAGAAGATTAGCTGAcgccaggcggccgcttggccggatccaggggcggtgcagggcggccggccctgcctgcgg

To test compounds for NSD2 degradation, a compound plate was made by diluting compounds in DMSO with the cell media in a 384-deep-well plate (Revvity). Then, 6 µL of the diluted compounds were transferred into the well of the 384-well assay plate (Corning). 24 µL of the NSD2 HiBiT U2OS P112 cells or parental U2OS cells (3000 cells/well) were added to each well to make a final volume of 30 µL with a 0.1% final DMSO concentration. For multiple time points, multiple assay plates were made and incubated in the incubator at 37°C in 5% CO_2_ for 3/6/12/24/48 hrs. 10 minutes before the reading, the assay plate and the Nano-Glo HiBiT Lytic Detection System (Promega) were to the room temperature. The HiBiT lytic reagent master mix was prepared according to the product instructions and added to the assay plate. The plate was then incubated on an orbital shaker for 5 minutes at room temperature, equilibrated for 10 minutes, followed by reading the plate using the GloMax luminometer (Promega) for luminescence detection. Compound treatment was normalized to the DMSO control with the background subtraction using the parental U2OS cell line as the background control plated alongside the HiBiT U2OS P112 cells. The data and graphs were then analyzed and generated using GraphPad Prism (v 10.4.1). DC50 and Dmax were calculated by fitting the data to a non-linear regression curve.

### Cell Viability Assay

The cell viability of NSD2 HiBiT U2OS P112 and parental U2OS cell lines treated with NSD2 degrader was determined by the CellTiter-Glo 2.0 Cell Viability Assay (Promega, G9242). 384-well plates (Corning) with cells and the reagent were equilibrated to the room temperature before detection. The reagent was added to the assay plate according to the product instruction and allowed to incubate on an orbital shaker for 3 minutes at room temperature before subjecting the plate to the GloMax luminometer (Promega). The data and graphs were then analyzed and generated using GraphPad Prism (v 10.4.1)

### Differential Scanning Fluorimetry

9.5 µL of the master mix containing FBXO22 FIST WT or variants, SYPRO Orange Dye (Invitrogen), and DSF buffer (20 mM Tris 7.5, 150 mM NaCl, and 1 mM TCEP) were added to a 384-well qPCR plate (Genesee Scientific). 0.5 µL of the compound in DMSO was added to make a final concentration of 15 µM of the protein and 15x SYPRO Orange. The DSF assay plate was covered and incubated on an orbital shaker at room temperature for 30 minutes. The DSF assay was performed using the ViiA 7 Real-Time PCR System (Applied Biosystems). The plate was heated from 25°C to 95°C with a rate of 0.033°C/second. The raw fluorescence value was analyzed using the Protein Thermal Shift Software (Applied Biosystems, v1.4) to obtain the T_m_. The graph was generated using GraphPad Prism (v 10.4.1).

### Cell-based Protein Degradation Assays

HEK293T and U2OS were obtained from ATCC and cultured in high glucose containing Dulbecco’s Modified Eagle’s Medium (DMEM; Gibco; cat. #11995) supplemented with 10% fetal bovine serum (VWR) and 1% penicillin/streptomycin (Gibco). Cells were treated with UNC10415667, UNC12149, hemin (Sigma; cat. #H9039), or DMSO control for the indicates times and analyzed by western blot using standard procedures.

Briefly, cells were lysed on ice for 15 minutes in NETN lysis buffer (20 mM Tris pH 8.0, 100 mM NaCl, 0.5 mM EDTA, 0.5% NP40) supplemented with 10 µg/mL aprotinin, 10 µg/mL leupeptin, 10 µg/mL pepstatin A, 1 mM sodium orthovanadate, 1 mM sodium fluoride, and 1 mM AEBSF (4-[two aminoethyl] benzenesulfonyl fluoride). Cell lysates were centrifuged at 20,000×g in a benchtop microcentrifuge at 4°C for 10 minutes.

Protein concentration was determined by Bradford assay (Bio-Rad; cat. #5000006). Protein concentration was normalized using Laemmli buffer. Samples were heated for 5 minutes at 95° in a dry water-bath. Samples were separated on TGX stain free gels (Bio-Rad) transferred to nitrocellulose membrane and blocked in 5% non-fat dry milk (Biorad; cat. #1706404) diluted in 1x TBS-T (137 mM NaCl, 2.7 mM KCl, 25 mM Tris pH 7.6, 1% Tween-20). Primary antibody incubations (NSD2 (Abcam; ab75359), BACH1 (Bethyl; A303-058A), and GAPDH (Santa Cruz; sc-47724) were performed overnight at 4°C. Secondary antibodies (Peroxidase AffiniPure Goat Anti-Rabbit IgG (H+L) (Jackson Immuno; 111-035-003) and Peroxidase AffiniPure Goat Anti-Mouse IgG (H+L) (Jackson Immuno; 115-035-003)) were incubated for one hour at room temperature. Blots were developed by chemiluminescence using Pierce ECL (Thermo Fisher Scientific; cat. #32106).

| Supplementary Table 1 | SCF-FBXO22, NSD2, UNC10088 | SCF-FBXO22, NSD2, UNC10415667 |
| --- | --- | --- |
| EMDB ID | (EMDB-xxxx) | (EMD-xxxx) |
| PDB ID | (PDB xxxx) | (PDB-xxxx) |
| Data collection and processing |  |  |
| Magnification | 130k x |  |
| Voltage (kV) | 300 |  |
| Electron exposure (e-/Å^2) | 40 |  |
| Detector | Thermo Fisher Falcon 4i |  |
| Defocus range (µm) | -1.0 to -2.0 µm (0.2 increments) |  |
| Pixel size (Å) | 0.951 |  |
| Symmetry imposed | C1 |  |
| Initial particle images (no.) | 4,324,385 | 11,321,391 |
| Final particle images (no.) | 84,420 | 47,442 |
| Map sharpening B factor (Å^2) | n/a | n/a |
| Map resolution (Å) | 3.8 | 5.7 |
| FSC threshold | 0.143 |  |
| Map resolution range (Å) | 3.6-4.3 | 4.6-5.9 |
| Model Refinement |  |  |
| Initial model used | AF3, PDB-6UE6 | AF3, PDB-6UE6 |
| Model resolution (Å) | 4.3 | 7.0 |
| FSC threshold | 0.5 |  |
| Model Composition |  |  |
| Non-hydrogen atoms | 12,384 | 12,388 |
| Protein residues | 1540 | 1540 |
| Ligands | 1 | 1 |
| B Factors |  |  |
| Protein (Å^2) min/max/mean | 78.0/362.9/163.2 | 120.3/669.2/273.6 |
| Ligand (Å^2) | 171.0 | 320.4 |
| R.M.S. Deviations |  |  |
| Bond lengths (Å) | 0.004 | 0.005 |
| Bond angles (°) | 0.932 | 1.189 |
| Validation |  |  |
| MolProbity score | 1.93 | 2.18 |
| Clashscore | 9.77 | 16.79 |
| Poor rotamers (%) | 0.07 | 0.00 |
| Ramachandran plot |  |  |
| Favored (%) | 93.63 | 92.84 |
| Allowed (%) | 6.37 | 7.16 |
| Disallowed (%) | 0.00 | 0.00 |

